# Visual detection of binary, ternary, and quaternary protein-protein interactions in fission yeast by Pil1 co-tethering assay

**DOI:** 10.1101/2021.04.13.439745

**Authors:** Zhong-Qiu Yu, Xiao-Man Liu, Dan Zhao, Dan-Dan Xu, Li-Lin Du

**Affiliations:** National Institute of Biological Sciences, 102206 Beijing, China; Tsinghua Institute of Multidisciplinary Biomedical Research, Tsinghua University, 102206 Beijing, China

**Keywords:** protein-protein interactions, Pil1 co-tethering assay, binary, ternary, quaternary, fission yeast.

## Abstract

Protein-protein interactions are vital for executing nearly all cellular processes. To facilitate the detection of protein-protein interactions in living cells of the fission yeast *Schizosaccharomyces pombe*, here we present an efficient and convenient method termed the Pil1 co-tethering assay. In its basic form, we tether a bait protein to mCherry-tagged Pil1, which forms cortical filamentary structures, and examine whether a GFP-tagged prey protein colocalizes with the bait. We demonstrate that this assay is capable of detecting pairwise protein-protein interactions of cytosolic proteins, transmembrane proteins, and nuclear proteins. Furthermore, we show that this assay can be used for detecting not only binary protein-protein interactions, but also ternary and quaternary protein-protein interactions. Using this assay, we systematically characterized the protein-protein interactions in the Atg1 complex and in the phosphatidylinositol 3-kinase (PtdIns3K) complexes and found that Atg38 is incorporated into the PtdIns3K complex I via an Atg38-Vps34 interaction. Our data show that this assay is a useful and versatile tool and should be added to the routine toolbox of fission yeast researchers.

## Introduction

Protein-protein interactions play crucial roles in regulating and executing most cellular functions (Alberts, 1998; Gavin and Superti-Furga, 2003). The detection of whether two proteins are interacting partners can provide significant insights into understanding the cellular roles of proteins. To analyze pairwise protein-protein interactions, a variety of *in vitro* and *in vivo* methods have been developed. The *in vitro* methods like coimmunoprecipitation and pull-down examine the interactions outside of a living organism, thus may fail to detect protein-protein interactions that are sensitive to environments. By contrast, the *in vivo* methods allow studies of protein-protein interactions in the cellular context. The most popular *in vivo* method to study protein-protein interactions is the yeast two-hybrid (Y2H) system, in which the budding yeast *Saccharomyces cerevisiae* is used as a living test tube and protein-protein interactions are detected by the activation of reporter genes through the reconstitution of a transcriptional activator in the nucleus (Fields and Song, 1989). The drawbacks of the Y2H assay include self-activation when using certain proteins as bait and false negative results for proteins unable to enter the nucleus of budding yeast and proteins only exhibit interactions in their native organisms but not in budding yeast.

*In vivo* methods that can be applied in the native organisms include fluorescence-based methods such as fluorescence resonance energy transfer (FRET) (Truong and Ikura, 2001) and bimolecular fluorescence complementation (BiFC) (Kerppola, 2006). These methods enable direct visualization of protein-protein interactions in living cells of the native organisms. However, both FRET and BiFC have their own drawbacks, with the former requiring specialized equipment and yielding a low signal output, and the latter suffering from the irreversibility of the binding of the split fluorescent protein fragments and the tendency of the split fluorescent protein fragments to fold together spontaneously.

In addition to binary protein-protein interactions, proteins also engage in ternary, quaternary, and even higher order interactions (Alberts, 1998). To detect and characterize ternary protein-protein interactions in living cells, yeast three-hybrid (Y3H) system (Zhang and Lautar, 1996), three-chromophore FRET (3-FRET) (Galperin et al., 2004), multicolor BiFC (Hu and Kerppola, 2003), and BiFC-based FRET (Shyu et al., 2008) have been developed based on the Y2H, FRET, and BiFC methods. However, they suffer similar limitations as the corresponding original methods.

The fission yeast *Schizosaccharomyces pombe* is a widely-used and powerful model organism for dissecting the mechanisms of a diverse range of cellular processes (Hoffman et al., 2015). For example, in recent years, we and others have used *S. pombe* to study autophagy (Fukuda et al., 2020; Liu et al., 2018; Matsuhara and Yamamoto, 2016; Mukaiyama et al., 2009; Nanji et al., 2017; Pan et al., 2020; Sun et al., 2013; Suzuki et al., 2015; Yu et al., 2020; Zhao et al., 2016, 2020). To facilitate the detection of *in vivo* protein-protein interactions in fission yeast, we have developed an imaging-based assay termed the Pil1 co-tethering assay. By fusing bait proteins to mCherry-tagged Pil1, which localizes to distinctive filamentary structures (Kabeche et al., 2011), and fusing prey proteins to a GFP, YFP, or CFP tag, protein-protein interactions can be visually detected as the colocalization of fluorescence signals in living fission yeast cells. We found that this assay is widely applicable in detecting pairwise protein-protein interactions of cytosolic proteins, transmembrane proteins, and nuclear proteins. Moreover, with this assay, we systematically examined the binary interactions among subunits of the Atg1 complex and the binary, ternary, and quaternary interactions among subunits of two PtdIns3K complexes. These application cases demonstrate the usefulness of this assay.

## Results

### Basic design of the Pil1 co-tethering assay

Pil1 forms cortical filaments in fission yeast cells (Kabeche et al., 2011) (Fig. 1A). The distinctive localization pattern of Pil1 makes it an ideal anchor for imaging-based detection of protein-protein interactions. In our basic design of the Pil1 co-tethering assay, two plasmids are constructed and introduced into fission yeast cells. One plasmid ectopically expresses from a medium-strength promoter (the *41nmt1* promoter) a fusion between Pil1, the red fluorescent protein mCherry, and a bait protein. The other plasmid ectopically expresses from the *41nmt1* promoter a fusion between the green fluorescent protein GFP and a prey protein. If the prey protein interacts with the bait protein, the GFP signal colocalizes with the mCherry signal on filamentary structures (Fig. 1B).

**Fig. 1.**
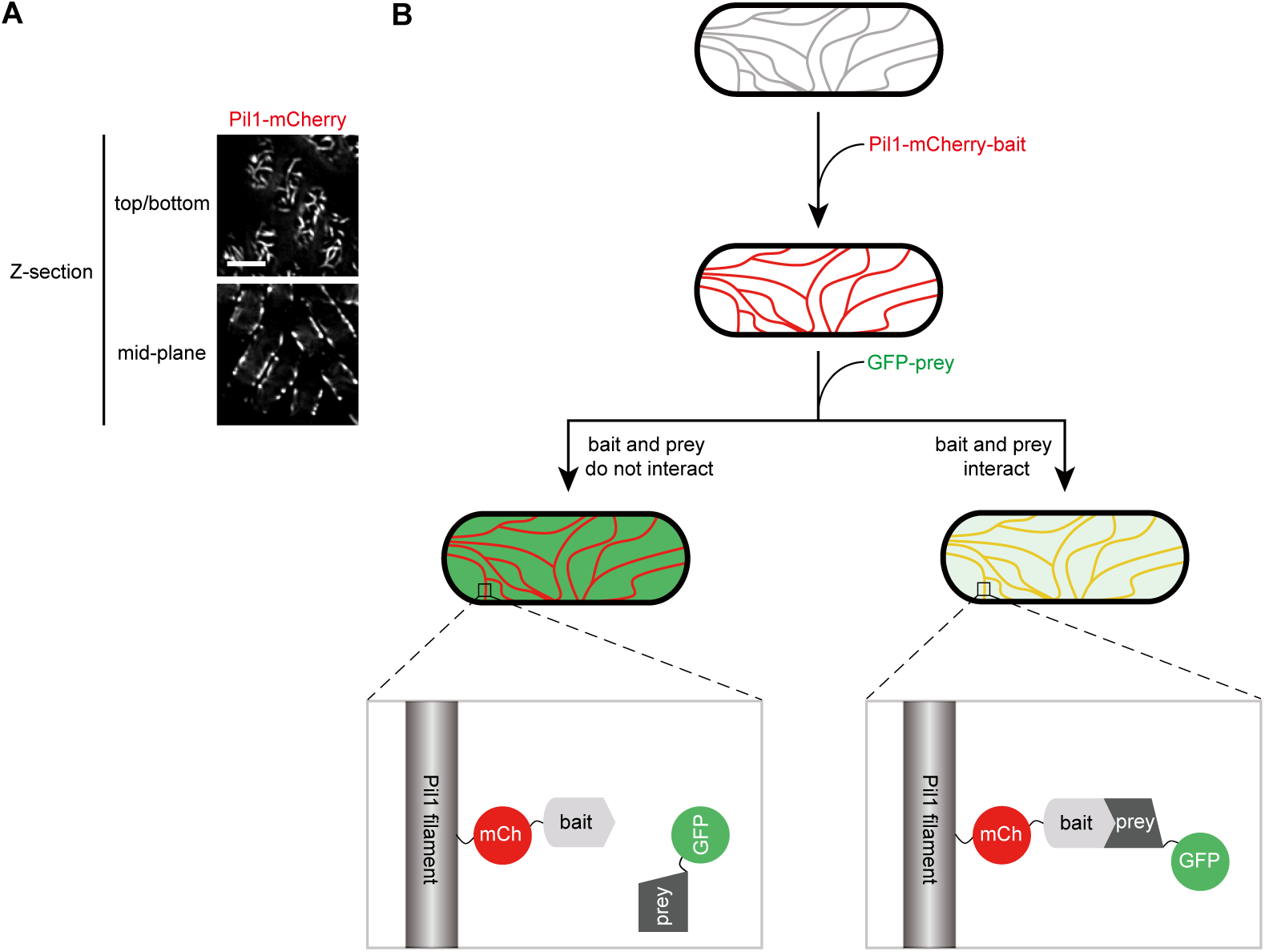
Basic design of the Pil1 co-tethering assay. (A) Localization of Pil1-mCherry in fission yeast. Images shown are deconvolved single optical sections, with one focused on the top (or bottom) of the cells, and the other focused in the mid-plane of the cells. Scale bar, 5 µm. A schematic of how the Pil1 co-tethering assay detects the interaction between bait and prey. The Pil1-mCherry-fused bait protein localizes to the Pil1 filaments in the cell cortex. If the GFP-fused prey protein interacts with the bait protein, the GFP signal colocalizes with the mCherry signal on the Pil1 filaments.

### Detecting the interactions between Atg8 and Atg8-interacting proteins using the Pil1 co-tethering assay

We first tested whether the Pil1 co-tethering assay can detect a previously reported interaction between two autophagy proteins, Atg8 and Atg38 (Yu et al., 2020). The ubiquitin-like protein Atg8 interacts with selective autophagy receptors and core autophagy-related (Atg) proteins via their Atg8-family-interacting motifs (AIMs) (Noda et al., 2010). Atg38 is a subunit of the PtdIns3K complex I (Araki et al., 2013; Yu et al., 2020). Fission yeast Atg38 contains an AIM (Yu et al., 2020) (Fig. 2A). We constructed a plasmid expressing a bait fusion protein consisting of Pil1 followed by mCherry and a 30-amino-acid Atg38 fragment, Atg38(161-190), which encompasses the AIM. This fusion protein localized to filament-like structures, in a manner similar to the distribution of Pil1-mCherry (Fig. 2B), suggesting that this fusion protein localizes to the Pil1 filaments. In cells expressing both Pil1-mCherry-Atg38(161-190) and GFP-tagged Atg8, the fluorescence signals of mCherry and GFP colocalized on the filamentary structures. As a negative control, in cells co-expressing Pil1-mCherry and GFP-Atg8, GFP- Atg8 showed a diffuse distribution in the cytosol and nucleus. Furthermore, mutating one or both of the two key residues in the AIM of Atg38, Phe178 and Val181, to alanine(s) totally abolished the colocalization on the filamentary structures. Mutating Pro52 and Arg67 in the AIM-binding region of Atg8 to alanines diminished the colocalization (Fig. 2B). These results are consistent with published results obtained using Y2H, coimmunoprecipitation, and pull-down assays (Yu et al., 2020).

**Fig. 2.**
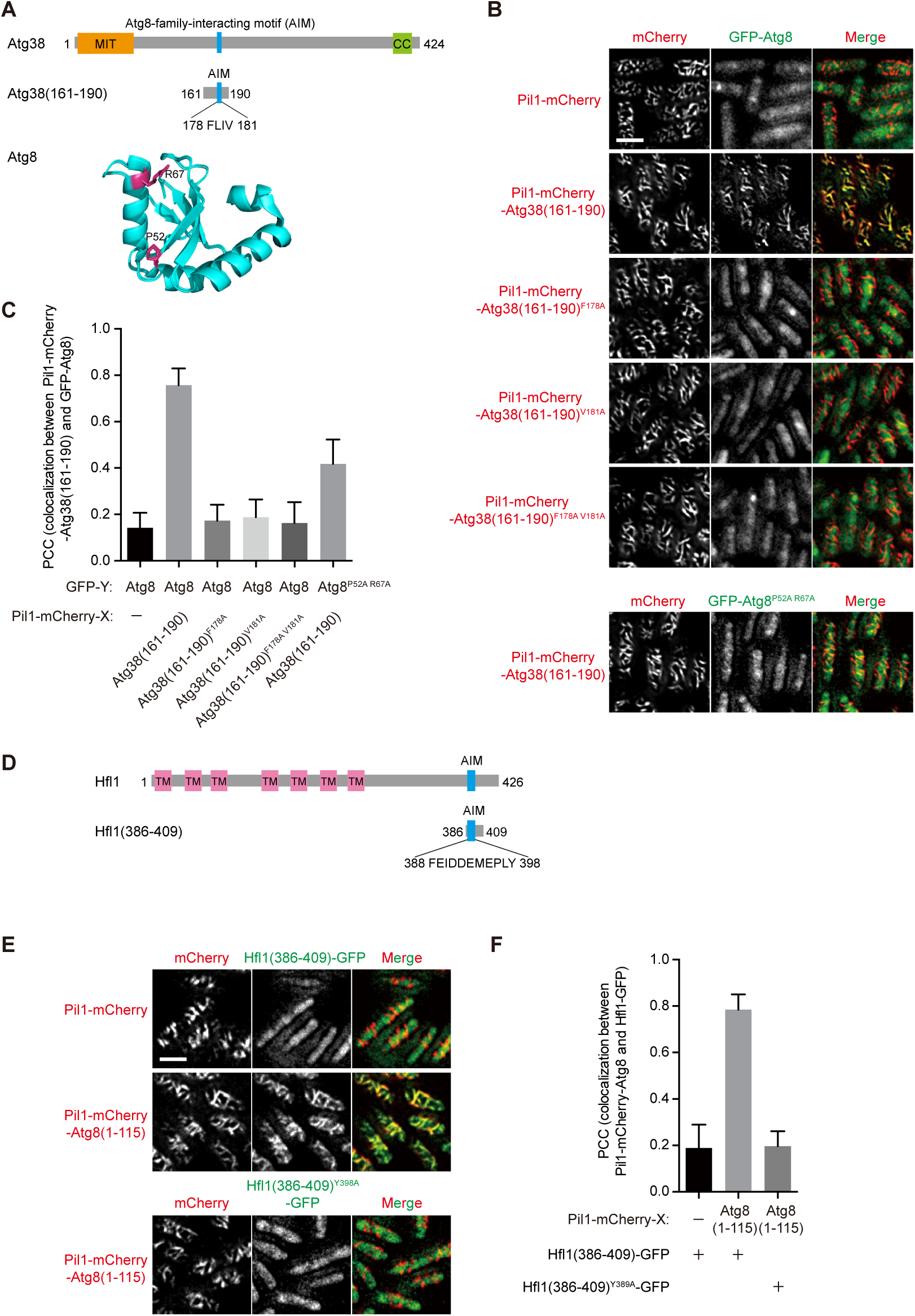
Detection of interactions between Atg8 and Atg8-interacting proteins using the Pil1 co-tethering assay. (A) Domain organization of Atg38 and the structure of Atg8. MIT, microtubule interacting and trafficking domain. CC, coiled-coil domain. AIM, Atg8-family- interacting motif. The structure of Atg8 (PDB 6AAF, chain A) is shown as a ribbon diagram with Pro52 and Arg67 highlighted in pink. (B) Atg38(161-190) interacts with Atg8 in the Pil1 co-tethering assay, and this interaction is blocked by the AIM mutations in Atg38(161-190) and diminished by the AIM-binding region mutation in Atg8. (C) Imaging data from the experiments shown in (B) were analyzed and the PCC values are presented as mean ± s.d. (10 cells). (D) Domain organization of Hfl1. TM, transmembrane domain. (E) Atg8 interacts with Hfl1(386-409) in the Pil1 co-tethering assay, and this interaction is blocked by the AIM mutation in Hfl1(386-409). (F) Imaging data from the experiments shown in (E) were analyzed and the PCC values are presented as mean ± s.d. (10 cells). Scale bars, 5 µm.

To quantitate the degree of colocalization between mCherry and GFP signals, we computed a Pearson correlation coefficient (PCC), whose values range from −1 to 1. Strong colocalization corresponds to a PCC value close to 1, whereas lack of colocalization corresponds to a PCC value close to 0 (Adler and Parmryd, 2010; Dunn et al., 2011). Consistent with the visual impression, the PCC values for the pairs of free Pil1 and Atg8 (negative control), Atg38(161-190) and Atg8, Atg38(161-190)^F178A^ and Atg8, Atg38(161-190)^V181A^ and Atg8, Atg38(161-190)^F178A V181A^ and Atg8, and Atg38(161-190) and Atg8^P52A R67A^ were 0.14, 0.76, 0.17, 0.19, 0.16, and 0.42, respectively (Fig. 2C). Thus, PCC values are useful quantitative measures of the pairwise interactions detected by the Pil1 co-tethering assay.

Next, we used the Pil1 co-tethering assay to examine a previously reported interaction between Atg8 and Hfl1 (Liu et al., 2018). Hfl1 is a vacuole membrane-localized protein containing seven transmembrane helices in its N-terminus and a noncanonical helical AIM in its C-terminal cytosolic tail (Fig. 2D). We used Atg8(1-115), which lacks the last six residues of Atg8, as bait, and a soluble fragment of Hfl1, Hfl1(386-409), which was previously shown to be sufficient for binding Atg8 (Liu et al., 2018), as prey. Hfl1(386-409)-GFP colocalized with Pil1-mCherry-Atg8(1-115) and this colocalization was abolished when the key residue in the helical AIM of Hfl1, Tyr398, was mutated to alanine (Fig. 2E,F). Together, these results obtained using Atg8 and its two binding proteins demonstrate that the Pil1 co-tethering assay is suitable to study interactions between cytosolic proteins in fission yeast.

### Detecting the interactions between transmembrane proteins using the Pil1 co-tethering assay

Interactions between transmembrane proteins are in general more difficult to detect than interactions between soluble proteins. Atg9 and Ctl1 are two multi-transmembrane proteins involved in autophagy and are known to interact with each other (Sun et al., 2013). Atg9 possesses four transmembrane helices and two reentrant membrane helices (Guardia et al., 2020; K et al., 2020; Maeda et al., 2020), and Ctl1 contains ten predicted transmembrane helices (Fig. 3A). We tested whether the Pil1 co-tethering assay can detect the interaction between Ctl1 and Atg9. Pil1-mCherry-Ctl1 showed a localization typical of the Pil1 filaments and the co-expressed Atg9-YFP localized to these filamentary structures (Fig. 3B,C). Thus, the Pil1 co-tethering assay can be used to study the interactions between transmembrane proteins.

**Fig. 3.**
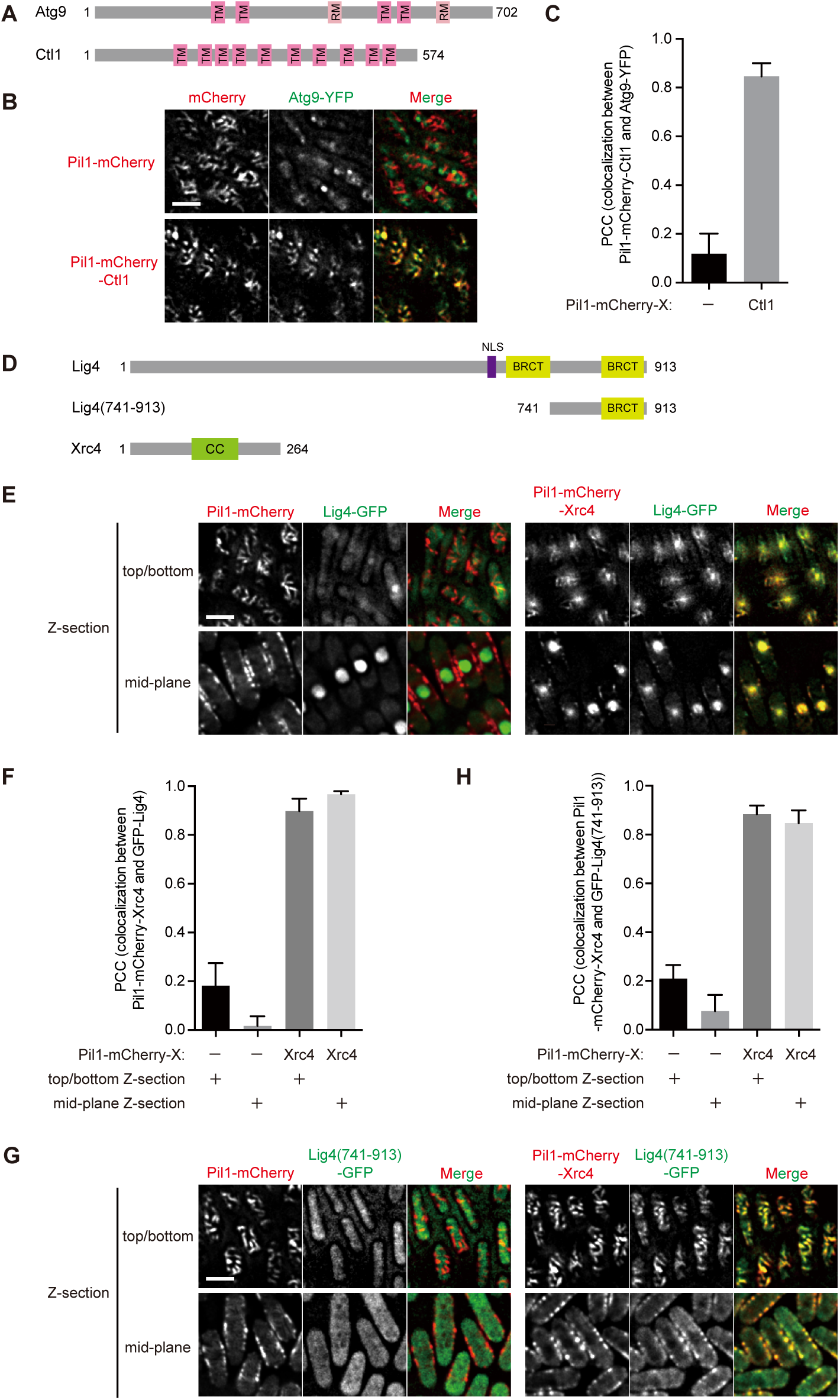
Detection of interactions between transmembrane proteins and interactions between nuclear proteins using the Pil1 co-tethering assay. (A) Domain organization of Atg9 and Ctl1. TM, transmembrane domain. RM, reentrant membrane domain. (B) Ctl1 interacts with Atg9 in the Pil1 co-tethering assay. (C) Imaging data from the experiments shown in (B) were analyzed and the PCC values are presented as mean ± s.d. (10 cells). (D) Domain organization of Lig4 and Xrc4. NLS, nuclear localization signal. BRCT, BRCT domain. CC, coiled-coil domain. (E) Xrc4 interacts with Lig4 in the Pil1 co-tethering assay. (F) Imaging data from the experiments shown in (E) were analyzed and the PCC values are presented as mean ± s.d. (10 cells). (G) Xrc4 interacts with Lig4(741-913) in the Pil1 co-tethering assay. (H) Imaging data from the experiments shown in (G) were analyzed and the PCC values are presented as mean ± s.d. (10 cells). Scale bars, 5 µm.

### Detecting the interactions between nuclear proteins using the Pil1 co-tethering assay

Pil1 filaments are cytoplasmic structures located outside of the nucleus. Therefore, we anticipated that the Pil1 co-tethering assay may encounter difficulty detecting interactions between nuclear-localized proteins. Nevertheless, we tested the Pil1 co-tethering assay using two nuclear-localized proteins Xrc4 and Lig4, which interact with each other and participate in the nonhomologous end joining (NHEJ) pathway of DNA double-strand break repair (Li et al., 2014). We chose Xrc4, which lacks a nuclear localization signal (NLS) and relies on Lig4 for its nuclear localization (Li et al., 2014) (Fig. 3D), as bait. As negative control, in cells expressing Pil1-mCherry, Lig4-GFP predominantly localized inside the nucleus (Fig. 3E). In contrast, in cells expressing Pil1-mCherry-Xrc4, a notable portion of Lig4-GFP colocalized with Pil1-mCherry-Xrc4 on cytoplasmic filamentary structures (Fig. 3E,F). Interestingly, when co-expressed with Lig4-GFP, a fraction of Pil1-mCherry-Xrc4 localized to the nucleus (Fig. 3E), presumably due to the interaction with the fraction of Lig4-GFP localized in the nucleus, because when co-expressed with a truncated Lig4 fragment that lacks the NLS but is still capable of binding Xrc4 (Li et al., 2014) (Fig. 3G,H), Pil1-mCherry-Xrc4 no longer exhibited the nucleus-localized signals (Fig. 3G). These results indicate that it is feasible to use the Pil1 co-tethering assay to study interactions between nuclear proteins, as NLS-mediated nuclear targeting does not completely prevent Pil1-fused bait and its binding partner from localizing to cytoplasmic filaments.

### Systematically probing the interactions among subunits of the Atg1 complex using the Pil1 co-tethering assay

The fission yeast Atg1 complex plays important roles in the initiation of starvation-induced autophagy and includes five components, namely, Atg1, Atg11, Atg13, Atg17, and Atg101 (Nanji et al., 2017; Pan et al., 2020; Sun et al., 2013; Suzuki et al., 2015) (Fig. 4A). We applied the Pil1 co-tethering assay to exhaustively examined the pairwise interactions among the five subunits, including self-interactions. To present the results in a concise manner, we used the PCC values to classify the results into three categories: strong colocalization, weak colocalization, and no obvious colocalization. Strong colocalization corresponds to PCC values greater than 0.7. Weak colocalization corresponds to PCC values less than 0.7 but greater than a threshold value. No obvious colocalization corresponds to PCC values less than the threshold value. The threshold value is either 0.3 or the PCC value obtained using free GFP as prey plus 0.05, whichever number is greater. In most cases, this threshold value is 0.3 (Fig. 4C). The only instance that this threshold value is greater than 0.3 is when using Pil1-mCherry-Atg1 as bait. In that instance, the control using the free GFP prey yielded a PCC value of 0.37 and thus the threshold was set at 0.42 (Fig. S1C,D).

**Fig. 4.**
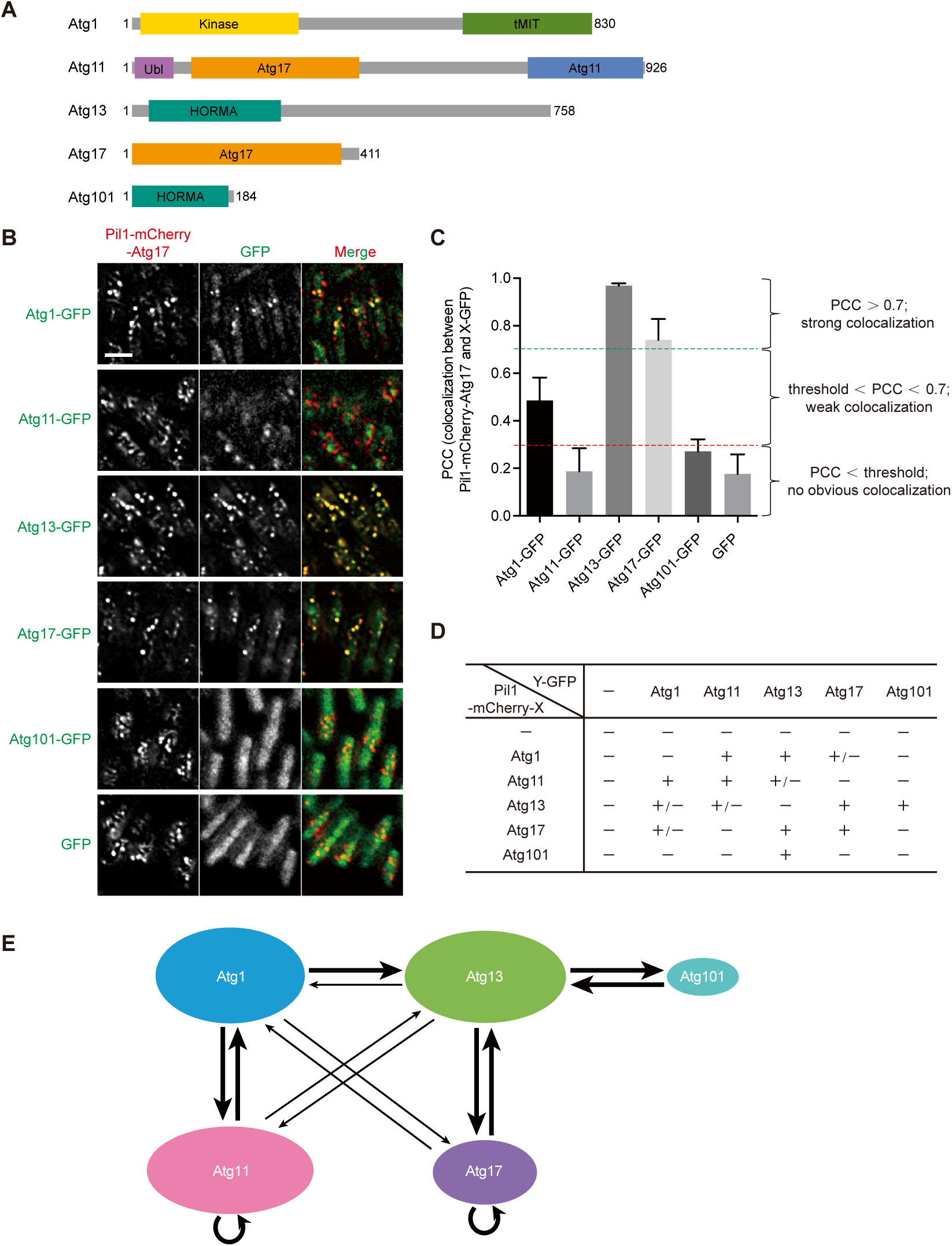
Mapping the interactions among subunits of the Atg1 complex using the Pil1 co- tethering assay. (A) Domain organization of subunits of the Atg1 complex. Kinase, kinase domain. tMIT, tandem MIT domain. Ubl, ubiquitin-like domain. Atg17, Atg17 domain. Atg11, Atg11 domain. HORMA, HORMA domain. CC, coiled-coil domain. (B) Detection of interactions between Atg17 and subunits of the Atg1 complex using the Pil1 co-tethering assay. Scale bar, 5 µm. (C) Imaging data from the experiments shown in (B) were analyzed and the PCC values are presented as mean ± s.d. (10 cells). (D) Summary of the interactions among subunits of the Atg1 complex revealed by the Pil1 co-tethering assay. “+” denotes a strong colocalization with the PCC value greater than 0.7. “+/−” denotes a weak colocalization with the PCC values less than 0.7 and greater than a threshold value, which is either 0.3 or the PCC value obtained using free GFP as prey plus 0.05, whichever number is greater. “−” denotes lack of colocalization with the PCC value less than 0.3. (E) A diagram of protein-protein interaction relationship among subunits of the Atg1 complex revealed by the Pil1 co-tethering assay. The arrow starts from a bait protein and points at a prey protein. The thick arrow denotes a strong colocalization, and the thin arrow denotes a weak colocalization.

We found bidirectional strong colocalizations (PCC > 0.7) between Atg1 and Atg11, between Atg13 and Atg17, and between Atg13 and Atg101 (Fig. S1, S2A,B,C,D, 4B,C,D,E), indicating that Atg1 tightly interacts with Atg11 and that Atg13 tightly interacts with Atg17 and Atg101. The Atg1-Atg13 pair exhibited strong colocalization in one direction and weak interaction in another direction (Fig. S1C,D, S2A,B, 4D,E). These four pairs of interactions are consistent with previously published results obtained using in-vitro pull-down of recombinant proteins (Nanji et al., 2017; Pan et al., 2020), indicating that these pairs of relatively strong interactions identified by the Pil1 co-tethering assay are direct physical interactions.

In addition to these relatively strong colocalizations, we detected weak colocalizations between Atg1 and Atg17 and between Atg11 and Atg13 (Fig. S1, S2A,B, 4B,C,D,E). The colocalization between Atg11 and Atg13 is independent of endogenous Atg1 (Fig. S2E,F,G,H), excluding the possibility that this colocalization is bridged by endogenous Atg1 through the Atg1-Atg11 interaction and the Atg1-Atg13 interaction. Thus, the interaction between Atg11 and Atg13 may be direct, despite being relatively weak in the Pil1 co-tethering assay.

Atg11 and Atg17 were also observed to self-interact in the Pil1 co-tethering assay (Fig. S1E,F, 4B,C,D,E), consistent with previously published results showing that both Atg11 and Atg17 can homodimerize (Nanji et al., 2017; Pan et al., 2020). Together, within the fission yeast Atg1 complex, the Pil1 co-tethering assay recapitulated six previously known binary interactions (Atg1-Atg11, Atg13-Atg17, Atg13-Atg101, Atg1-Atg13, Atg11-Atg11, and Atg17-Atg17), and identified two previously unknown binary interactions (Atg1-Atg17 and Atg11-Atg13). These results demonstrate the usefulness of the Pil1 co-tethering assay in detecting protein-protein interactions within a multiprotein complex.

### Characterizing the binary interactions among subunits of PtdIns3K complexes using the Pil1 co-tethering assay

In fission yeast, there are two PtdIns3K complexes: the PtdIns3K complex I, which functions in autophagy, and the PtdIns3K complex II, which participates in vacuolar protein sorting. These two complexes share three common subunits: Vps15, Vps34, and Atg6; complex I possesses two specific subunits: Atg14 and Atg38; complex II possesses one specific subunit: Vps38 (Yu et al., 2020) (Figure 5A). It remains incompletely understood how these two complexes are organized, and in particular, how Atg38 is integrated into the PtdIns3K complex I. To further our understanding of these two complexes, we applied the Pil1 co-tethering assay to systematically examine all pairwise combinations of the six proteins. We observed four pairs of bidirectional strong colocalizations (PCC > 0.7): Vps15 and Vps34, Vps34 and Atg38, Atg6 and Atg14, and Atg6 and Vps38 (Fig. S3, S4, S5, 5B,C). Additionally, weak colocalizations (0.3 < PCC < 0.7) were observed between Vps15 and Atg6, between Vps15 and Atg14, between Vps15 and Vps38, between Vps34 and Atg6, and between Atg6 bait and Atg6 prey (Fig. S4, S5, 5B,C). Thus, we obtained a protein-protein interaction map of the PtdIns3K complexes (Figure 5C). Among these interactions, only the interaction between Atg6 and Vps38 and the Atg6 self-interaction were detected in a proteome-wide Y2H analysis (Vo et al., 2016), suggesting that the Pil1 co-tethering assay has high sensitivity in detecting binary interactions within multiprotein complexes.

**Fig. 5.**
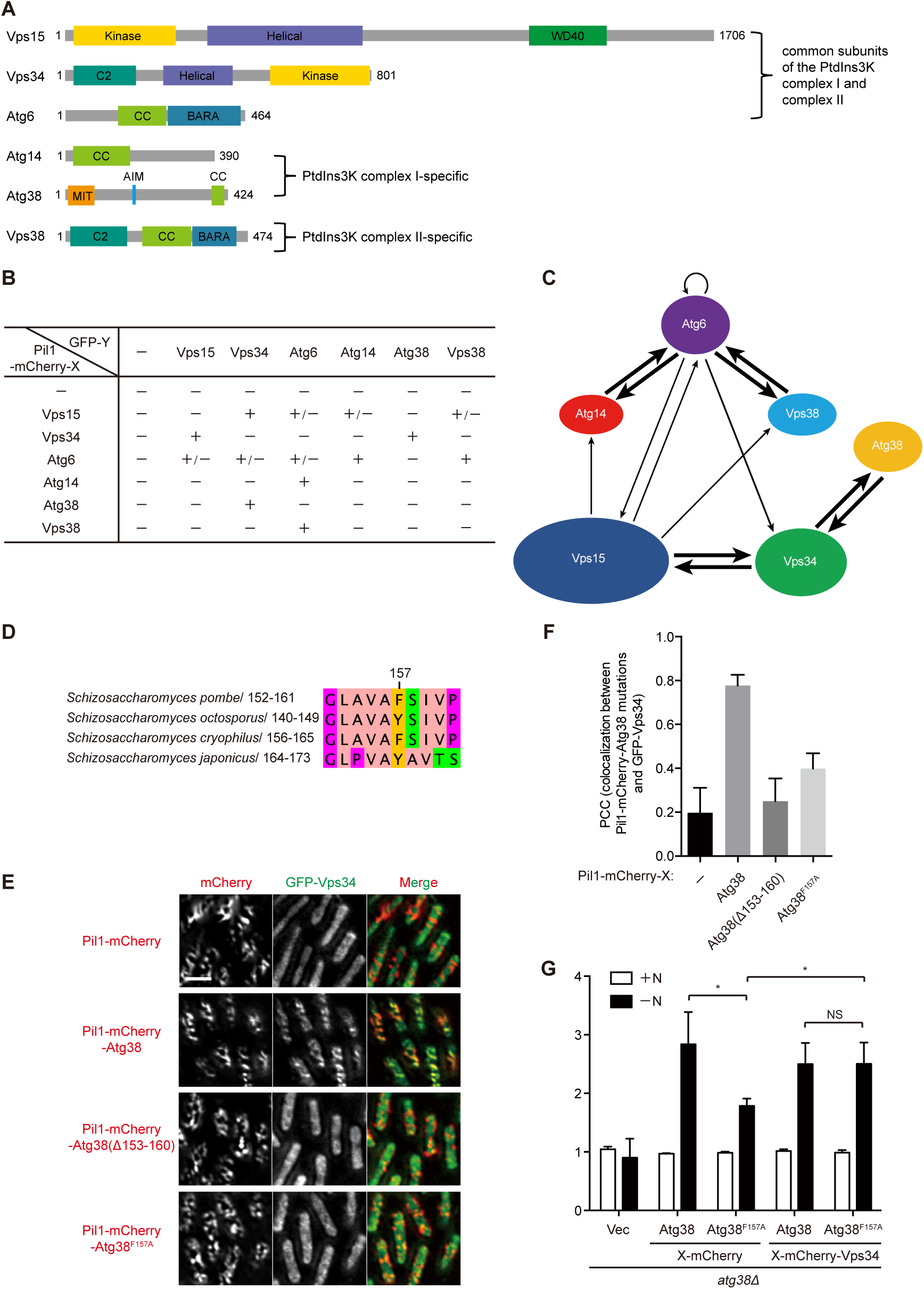
Mapping the interactions among subunits of PtdIns3K complexes using the Pil1 co-tethering assay. (A) Domain organization of subunits of the two PtdIns3K complexes. Kinase, kinase domain. Helical, helical domain. WD40, WD40 domain. C2, C2 domain. CC, coiled-coil domain. BARA, beta-alpha repeated, autophagy-specific domain. MIT, microtubule interacting and trafficking domain. (B) Summary of the interactions among subunits of PtdIns3K complexes revealed by the Pil1 co-tethering assay. “+” denotes a strong colocalization. “+/−” denotes a weak colocalization. “−” denotes no obvious colocalization. (C) A diagram of protein-protein interaction relationship among subunits of PtdIns3K complexes revealed by the Pil1 co-tethering assay. The arrow starts from a bait protein and points at a prey protein. The thick arrow denotes a strong colocalization, and the thin arrow denotes a weak colocalization. (D) A region of Atg38 conserved among *S. pombe*, *S. octosporus*, *S. cryophilus*, and *S. japonicus*. (E) F157 in Atg38 is important for its interaction with Vps34 in the Pil1 co- tethering assay. (F) Imaging data from the experiments shown in (E) were analyzed and the PCC values are presented as mean ± s.d. (10 cells). (G) Autophagic flux measurement using the Pho8Δ60 assay was performed in *atg38Δ* cells transformed with an empty vector or a plasmid expressing wild-type Atg38, Atg38^F157A^, wild-type Atg38 fused with Vps34, or Atg38^F157A^ fused with Vps34. Cells were collected before (+N) and after culturing in nitrogen- free medium for 4 h (−N). Average activity from non-starved (+N) samples was set to 1. Data are mean ± s.d. of triplicates from representative experiments. * indicates *P* < 0.05; NS, not significant. *P* values were calculated using Welch’s t-test. Scale bars, 5 µm.

To ascertain whether the four pairs of strong colocalizations detected by the Pil1 co-tethering assay reflect direct interactions or indirect interactions bridged by other subunits of the complexes, we performed the Pil1 co-tethering assay in cells lacking subunits other than those used as bait and prey. For the Vps15-Vps34 pair, we deleted the four genes encoding the other subunits of the two PtdIns3K complexes and found that in the absence of Atg6, Atg14, Atg38, and Vps38, the interaction between Vps15 and Vps34 remained unchanged (Fig. S6A,B,C,D). Similarly, we found that the interaction between Vps34 and Atg38 is independent of all the other subunits of the PI3K complex I, Vps15, Atg6, and Atg14 (Fig. S6E,F,G,H), the interaction between Atg6 and Atg14 is independent of all the other subunits of the PI3K complex I, Vps15, Vps34, and Atg38 (Fig. S6I,J,K,L), and the interaction between Atg6 and Vps38 is independent of all the other subunits of the PI3K complex II, Vps15 and Vps34 (Fig. S6M,N,O,P). These results suggest that the four pairs of strong protein-protein interactions are not mediated by any other subunits of the corresponding complex(es) and are probably direct interactions.

A 4.4 Å resolution crystal structure of the budding yeast PtdIns3K complex II has been reported (Rostislavleva et al., 2015). In that structure, the buried surface areas between Vps15 and Vps34, between Atg6 and Vps38, between Vps15 and Atg6, between Vps15 and Vps38, between Vps34 and Atg6, and between Vps34 and Vps38 are 3528 Å^2^, 2496 Å^2^, 921 Å^2^, 1702 Å^2^, 92 Å^2^, and 24 Å^2^, respectively. Given that binding affinity directly correlates with the amount of buried surface areas (Chen et al., 2013), the interactions between Vps15 and Vps34 and between Atg6 and Vps38 are likely stronger than the other pairs of interactions within the PtdIns3K complex II, consistent with our observations in the Pil1 co-tethering assay that Vps15 strongly colocalized with Vps34, and Atg6 strongly colocalized with Vps38. Because the low resolution electron microscopy structures of the PtdIns3K complex I showed that it adopts an overall structure similar in shape to that of the PtdIns3K complex II (Baskaran et al., 2014; Ma et al., 2017; Young et al., 2019), and the complex I-specific subunit Atg14 is known to bind to Atg6 in a manner similar to the complex II-specific subunit Vps38 (Itakura et al., 2008), it is likely that there is also a large buried surface area between Atg6 and Atg14 in the PtdIns3K complex I, consistent with the strong colocalization between Atg6 and Atg14 in the Pil1 co-tethering assay. The structural relationship between Atg38 and other subunits of PtdIns3K complex I has not been clearly resolved, but the published structural studies on budding yeast Atg38 and the human homolog of Atg38 (called NRBF2) do not support any extensive contact between Atg38/NRBF2 and Vps34 (Ohashi et al., 2016; Young et al., 2016, 2019). Thus, the four pairs of strong colocalizations we observed using the Pil1 co-tethering assay include three pairs (Vps15-Vps34, Atg6-Vps14, and Atg6-Vps38) that are consistent with previous structural knowledge and one pair (Vps34-Atg38) that is unexpected from previous structural knowledge obtained using budding yeast and human proteins, suggesting that the structural organization of the PtdIns3K complex I in fission yeast may be different from that in budding yeast and humans.

### The Vps34-Atg38 interaction is important for autophagy

In a sequence alignment of Atg38 proteins from four fission yeast species belonging to the *Schizosaccharomyces* genus, we noticed that, between the MIT domain and the AIM, there exists a conserved stretch of 10 amino acids located between residues 152 and 161 of *S. pombe* Atg38 (Fig. 5D). We wondered whether this region is required for the interaction between Vps34 and Atg38. Using the Pil1 co-tethering assay, we found that, either deleting residues 153-160, which are predicted to adopt a β strand conformation, or mutating Phe157 to alanine largely blocked the Vps34-Atg38 interaction (Fig. 5E,F). To further investigate whether this interaction is important for autophagy, we used the Pho8Δ60 assay, in which Pho8Δ60 is transported into the vacuole in an autophagy-dependent manner and activated by vacuolar proteases (Noda and Klionsky, 2008; Yu et al., 2020), to analyze the effect of the Atg38-F157A mutation on autophagy. This mutation diminished the starvation-induced increase of Pho8Δ60 activity, and fusing Vps34 to the F157A-mutated Atg38 rescued this impairment (Fig. 5G), indicating that the Vps34-Atg38 interaction is important for autophagy.

### Detecting the ternary Vps15-Vps34-Atg38 interaction using the Pil1 co-tethering assay

Because Vps34 tightly interacts with Vps15 and Atg38 and both binary interactions are independent of the other subunits of PtdIns3K complexes, we hypothesized that Vps34 may bridge the association between Vps15 and Atg38 in the assembly of the PtdIns3K complex I. To test this idea, we introduced into the Pil1 co-tethering assay system a third plasmid ectopically expressing from the *41nmt1* promoter a cyan fluorescent protein (CFP)-tagged prey protein so that ternary protein-protein interactions can be detected. When CFP-tagged Vps34 was co-expressed, GFP-Atg38 strongly colocalized with Pil1-mCherry-Vps15 on filamentary structures, whereas no colocalization was observed without the ectopic expression of Vps34 (Fig. 6A,B). Similarly, ectopic expression of CFP-Vps34 also led to the colocalization between GFP-Vps15 and Pil1-mCherry-Atg38 (Fig. 6C,D). These results support the idea that Vps34 can bridge the interaction between Vps15 and Atg38. Thus, the Pil1 co-tethering assay can be used to detect ternary interactions.

**Fig. 6.**
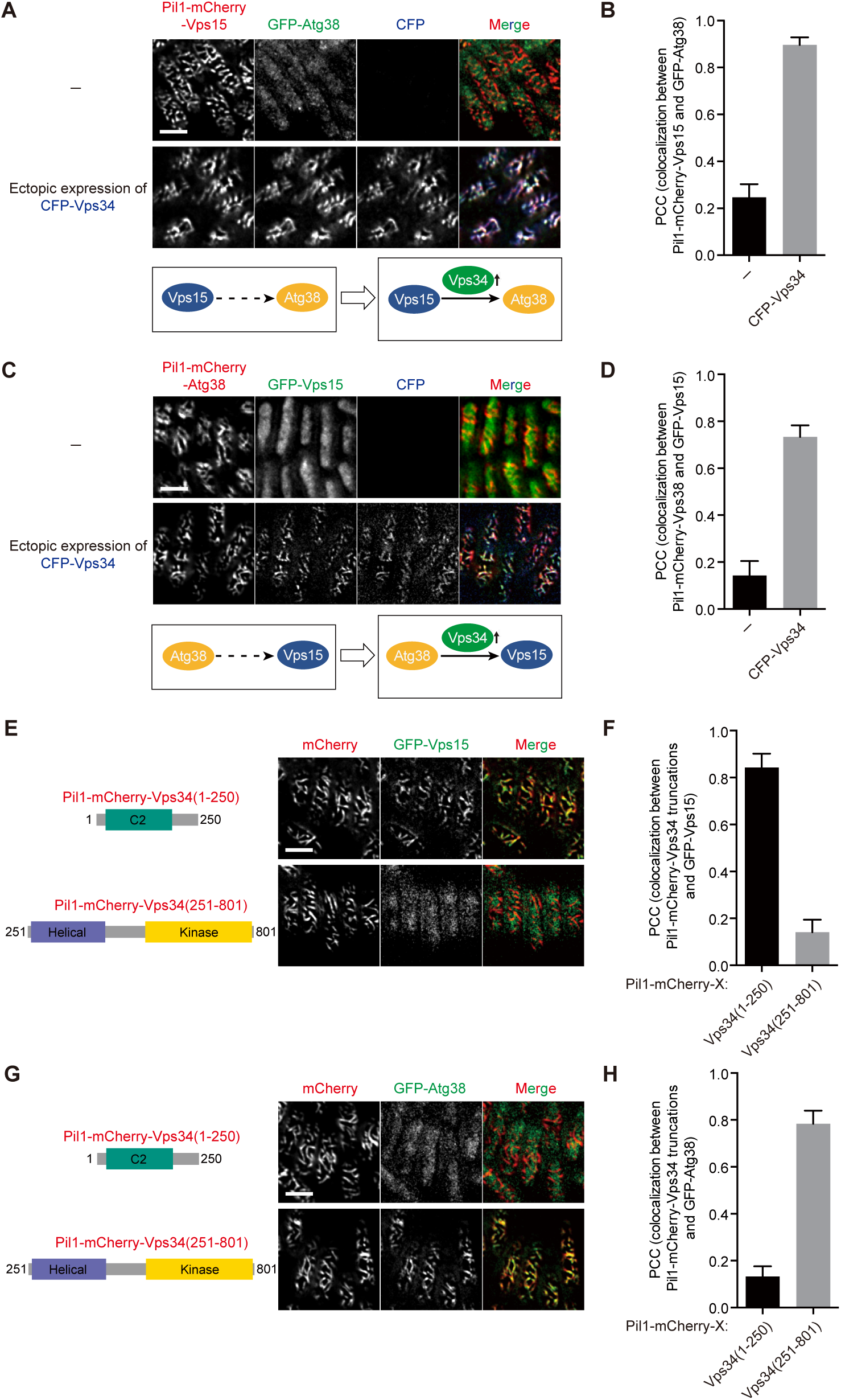
Analyzing the ternary Vps15-Vps34-Atg38 interaction using the Pil1 co-tethering assay. (A) Ectopic expression of Vps34 led to the colocalization of Pil1-mCherry-Vps15 and GFP-Atg38 in the Pil1 co-tethering assay. (B) Imaging data from the experiments shown in (A) were analyzed and the PCC values are presented as mean ± s.d. (10 cells). (C) Ectopic expression of Vps34 led to the colocalization of Pil1-mCherry-Atg38 and GFP-Vps15 in the Pil1 co-tethering assay. (D) Imaging data from the experiments shown in (C) were analyzed and the PCC values are presented as mean ± s.d. (10 cells). (E) Vps15 colocalized with the N- terminal region, but not the C-terminal region of Vps34 in the Pil1 co-tethering assay. (F) Imaging data from the experiments shown in (E) were analyzed and the PCC values are presented as mean ± s.d. (10 cells). (G) Atg38 colocalized with the C-terminal region, but not the N-terminal region of Vps34 in the Pil1 co-tethering assay. (H) Imaging data from the experiments shown in (G) were analyzed and the PCC values are presented as mean ± s.d. (10 cells). Scale bars, 5 µm.

Truncation analysis of Vps34 showed that residues 1-250 of Vps34 containing a C2 domain mediate its interaction with Vps15 (Fig. 6E,F), whereas residues 251-801 of Vps34 including a helical domain and a lipid kinase domain are responsible for binding Atg38 (Fig. 6G,H). Thus, Vps34 bridges the Vps15-Atg38 interaction by simultaneously binding both Vps15 and Atg38 through different regions.

We also examined whether ectopic expression of Vps34 promotes the interactions between Vps15 and the other three subunits of the two PtdIns3K complexes, Atg6, Atg14, and Vps38, which did not exhibit strong interactions with Vps34 in the binary Pil1 co-tethering assay. Unsurprisingly, none of them showed a stronger colocalization with Pil1-mCherry- Vps15 upon the ectopic expression of Vps34 (Fig. S7).

### Detecting the ternary interactions formed between Vps15 and either the Atg6-Atg14 subcomplex or the Atg6-Vps38 subcomplex

In the budding yeast *Saccharomyces cerevisiae*, Atg6 and Atg14 form a subcomplex in the PtdIns3K complex I and Atg6 and Vps38 form a subcomplex in the PtdIns3K complex II (Araki et al., 2013; Rostislavleva et al., 2015). Similarly, we observed using the Pil1 co-tethering assay that, in fission yeast, strong pairwise interactions exist between Atg6 and Atg14 and between Atg6 and Vps38 (Fig. S4, S5, 5B,C). However, none of these three proteins exhibited strong interactions with other subunits of the two PtdIns3K complexes (Fig. S3, S4, S5, 5B,C). We hypothesized that they may only strongly engage other subunits after forming the Atg6-Atg14 subcomplex or the Atg6-Vps38 subcomplex. To test this idea, we used the Pil1 co-tethering assay to examine whether ternary interactions exist between the Atg6-Atg14 subcomplex and the other subunits of the PtdIns3K complex I, and between the Atg6-Vps38 subcomplex and the other subunits of the PtdIns3K complex II.

In contrast to the observations that Vps15 showed no colocalization with Atg14 and Vps38 when using Atg14 and Vps38 individually as bait in the Pil1 co-tethering assay, upon ectopically expressing CFP-tagged Atg6, GFP-tagged Vps15 strongly colocalized with Pil1- mCherry-Atg14 as well as with Pil1-mCherry-Vps38 (Fig. 7A,B,C,D). Endogenous Vps34 is not required for these ternary interactions (Fig. 7A,B,C,D). When using Atg6 as bait, the ectopic expression of Atg14 or Vps38 notably enhanced the colocalization between Atg6 and Vps15 (Fig. 7E,F). These results suggest that the Atg6-Atg14 subcomplex and the Atg6-Vps38 subcomplex bind Vps15 more strongly than Atg6, Atg14, and Vps38 individually. Ectopic expression of Atg6 had no effect on the colocalizations between Atg14 and Vps34, between Vps38 and Vps34, and between Atg14 and Atg38 (Fig. S8). Taken together, these results suggest that the Atg6-Atg14 subcomplex and the Atg6-Vps38 subcomplex are respectively incorporated into the PtdIns3K complex I and the PtdIns3K complex II through engaging Vps15.

**Fig. 7.**
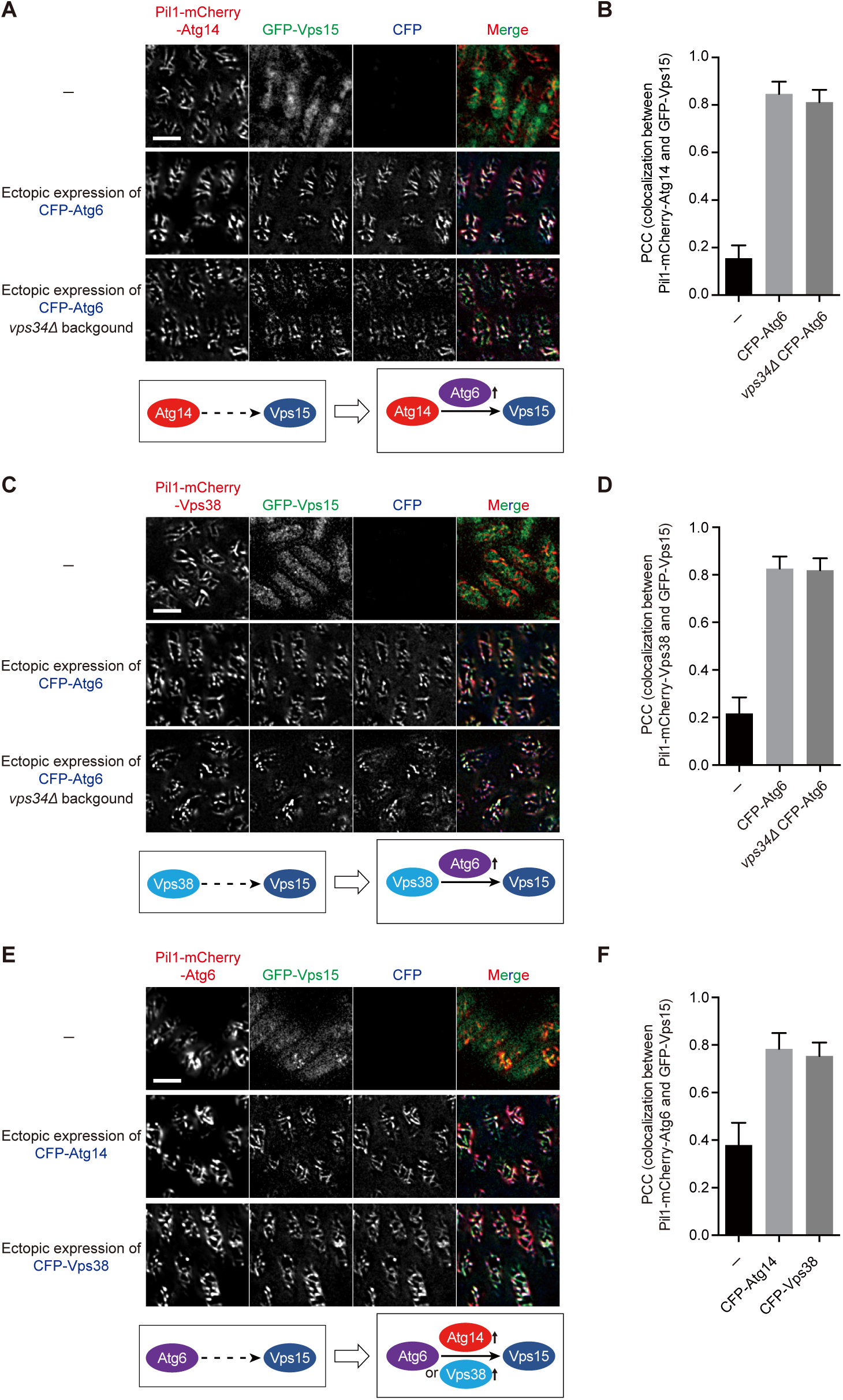
Analyzing the ternary Atg14-Atg6-Vps15 interaction and the ternary Atg14- Vps38-Vps15 interaction. (A) Ectopic expression of Atg6 led to the colocalization of Atg14 and Vps15, and this colocalization is independent of Vps34. (B) Imaging data from the experiments shown in (A) were analyzed and the PCC values are presented as mean ± s.d. (10 cells). (C) Ectopic expression of Atg6 led to the colocalization of Vps38 and Vps15, and this colocalization is independent of Vps34. (D) Imaging data from the experiments shown in (C) were analyzed and the PCC values are presented as mean ± s.d. (10 cells). (E) Ectopic expression of Atg14 or Vps38 enhanced the colocalization of Atg6 and Vps15. (F) Imaging data from the experiments shown in (E) were analyzed and the PCC values are presented as mean ± s.d. (10 cells). Scale bars, 5 µm.

### Detecting the quaternary interactions formed among Vps34, Vps15, and either the Atg6- Atg14 subcomplex or the Atg6-Vps38 subcomplex

Given that Vps34 strongly interacted with Vps15 and Vps15 strongly interacted with the Atg6-Atg14 subcomplex and the Atg6-Vps38 subcomplex, we hypothesized that Vps15 may bridge the association between Vps34 and these two subcomplexes. To test this, in the ternary Pil1 co-tethering assay systems using Pil1-mCherry-fused Vps34 as bait and CFP-tagged Atg6 and GFP-tagged Atg14 or Vps38 as preys, we ectopically expressed Vps15 by introducing a fourth plasmid expressing from the *41nmt1* promoter 13Myc-tagged Vps15. Without the ectopic expression of Vps15, no or little colocalizations were observed between preys and Pil1- mCherry-Vps34 (Fig. 8A,B,C,D). In contrast, when Vps15 was ectopically expressed, the preys obviously colocalized with Pil1-mCherry-Vps34 (Fig. 8A,B,C,D). Thus, using the Pil1 co-tethering assay we detected the quaternary Vps34-Vps15-Atg6-Atg14 interaction and the quaternary Vps34-Vps15-Atg6-Vps38 interaction.

**Fig. 8.**
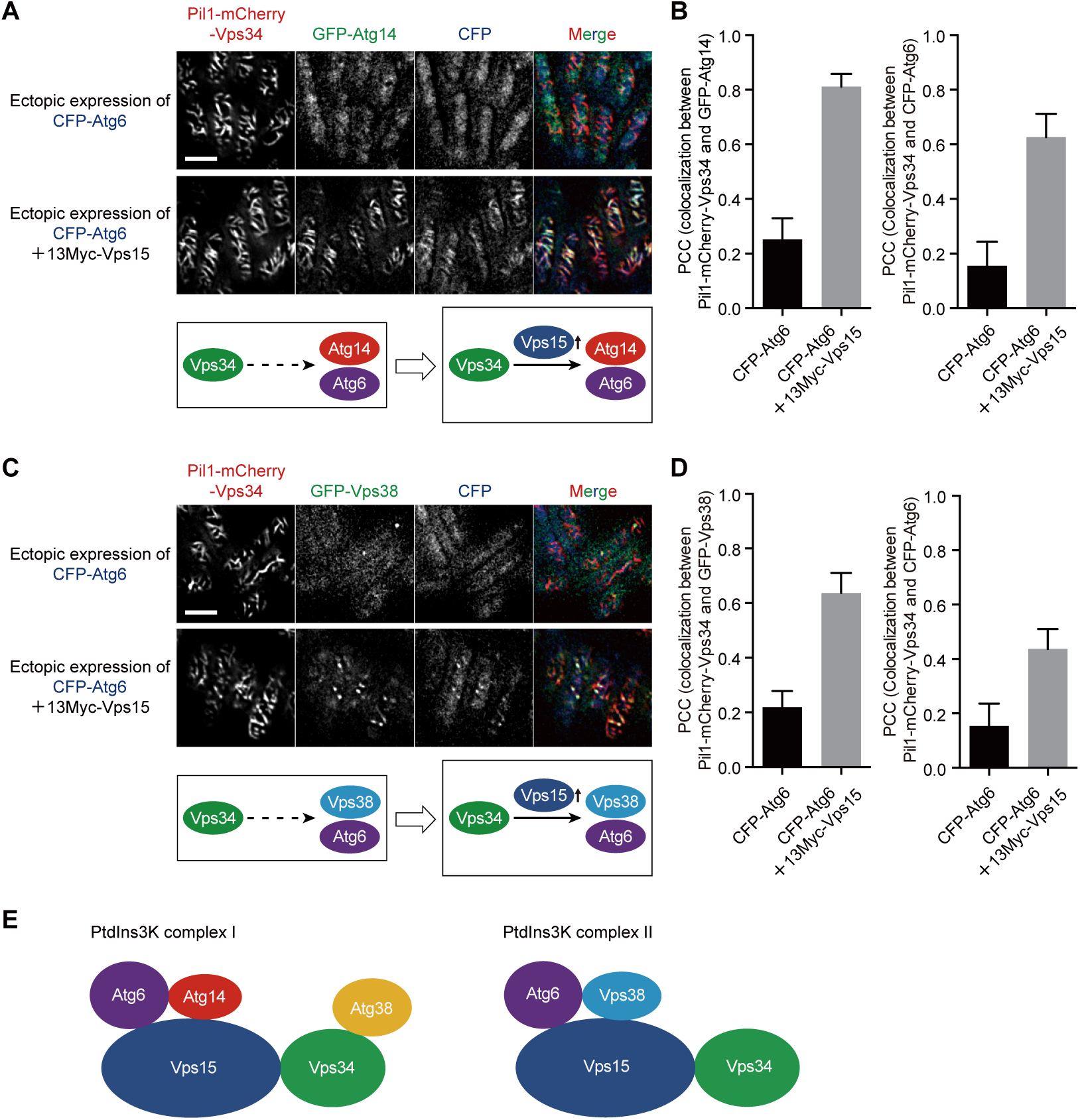
Vps15 bridges the interactions between Vps34 and the Atg14-Atg6 subcomplex and the interactions between Vps34 and the Vps38-Atg6 subcomplex. (A) Ectopic expression of Vps15 led to the colocalization of Vps34 and the Atg14-Atg6 pair in the Pil1 co-tethering assay. (B) Imaging data from the experiments shown in (A) were analyzed and the PCC values are presented as mean ± s.d. (10 cells). (C) Ectopic expression of Vps15 led to the colocalization of Vps34 and the Vps38-Atg6 pair in the Pil1 co-tethering assay. (D) Imaging data from the experiments shown in (C) were analyzed and the PCC values are presented as mean ± s.d. (10 cells). (E) Model of the organization of the PtdIns3K complex I and the PtdIns3K complex II in fission yeast. Scale bars, 5 µm.

Collectively, the binary, ternary, and quaternary interactions obtained using the Pil1 co-tethering assay revealed the organization of the two PtdIns3K complexes in fission yeast: in the PtdIns3K complex I, Vps15 bridges the association between the Atg6-Atg14 subcomplex and Vps34, which is the subunit linking Atg38 to the rest of the complex; in the PtdIns3K complex II, Vps15 bridges the association between the Atg6-Vps38 subcomplex and Vps34 (Fig. 8E).

## Discussion

Here we report a new method, which we termed the Pil1 co-tethering assay, to visually detect protein-protein interactions in fission yeast. In this method, the colocalization of GFP-, YFP-, or CFP-tagged prey protein(s) with a Pil1-mCherry-fused bait protein on visually distinctive cytoplasmic filaments indicates bait and prey proteins can interact with each other. The successful applications of this method to cytosolic proteins Atg8 and Atg8-interacting proteins, transmembrane proteins Atg9 and Ctl1, nuclear proteins Xrc4 and Lig4, components of the Atg1 complex, and components of the two PtdIns3K complexes demonstrate that the Pil1 co-tethering assay is an effective tool that can be broadly used to detect protein-protein interactions. In other organisms, imaging-based colocalization assays similar in principle but different in design have been used for the detection of binary protein-protein interactions (Blanchard et al., 2006; Gallego et al., 2013; Herce et al., 2013; Lv et al., 2017; Miller et al., 2007; Yurlova et al., 2014; Zolghadr et al., 2008). In this study, we expanded the applications of this class of assays to the detection of ternary and quaternary protein-protein interactions.

Using the Pil1 co-tethering assay, we detected two modes of ternary protein-protein interactions within the two PtdIns3K complexes. In the first mode, one protein bridges the interaction of two others, as exemplified by the ternary Vps15-Vps34-Atg38 interaction (Fig. 6A,B,C,D). In the second mode, a complex formed by two proteins, but not each protein individually, strongly interacts with the third protein, as in the scenarios of the Atg14-Atg6- Vps15 interaction and the Vps38-Atg6-Vps15 interaction (Fig. 7).

According to structural studies using proteins from budding yeast and humans, the two PtdIns3K complexes adopt a V-shaped architecture, in which Vps15 organizes these two complexes by bridging Vps34 and the Atg6-Atg14 subcomplex or the Atg6-Vps38 subcomplex (Baskaran et al., 2014; Ma et al., 2017; Rostislavleva et al., 2015). Our results of the binary, ternary, and quaternary interactions of Vps15, Vps34, Atg6, Atg14, and Vps38 (Fig. 5, 7, 8, S3, S4, S5, S6, S7, S8) support that the fission yeast PtdIns3K complexes share a similar overall structure with their counterparts in budding yeast and mammals. However, how the fifth subunit of the PtdIns3K complex I, Atg38 (NRBF2 in mammals), is incorporated into this complex seems vary among different species. Budding yeast Atg38 was initially reported to interact with Atg14 and Vps34 and thereby links the Vps15-Vps34 subcomplex and the Atg6- Atg14 subcomplex (Araki et al., 2013), but a later study failed to find evidence supporting the Atg38-Vps34 interaction (Ohashi et al., 2016). Mammalian NRBF2 was shown to interact with ATG14 and BECN1/Beclin 1 (homolog of yeasts Atg6) but not Vps34 (Young et al., 2016, 2019). Differing from the situations in budding yeast and mammals, we found that fission yeast Atg38 is incorporated into the PtdIns3K complex I by binding to the 251-801 region of Vps34 consisting of a helical domain and a lipid kinase domain (Fig. 6).

The newly established Pil1 co-tethering assay in fission yeast has the following advantages: (1) Compared to in vitro methods, this assay does not need to isolate proteins from their native cellular environments, thus can better preserve protein-protein interactions; (2) Compared to the popular Y2H assay, this assay does not need a reporter gene, thus avoiding false positives or false negatives associated with the use of reporter genes; (3) This assay does not require specialized equipment or technical expertise: fluorescent proteins-fused bait and prey proteins can be visualized using a regular fluorescent microscope and the interaction is reported by their colocalization on the Pil1 filaments in living cells, an easy and straightforward readout; (4) The introduction of PCC to evaluate the degree of colocalization between bait and prey proteins allows the strengths of the interactions to be measured in a quantitative manner; (5) This assay is useful for detecting and characterizing not only binary interactions, but also ternary and quaternary interactions.

On the other hand, like other protein-protein interaction assays, the Pil1 co-tethering assay has limitations. First, proteins that interact with Pil1 cannot be used in this assay. Such proteins should be rare and the use of the free Pil1 control should be able to identify such situations. Second, protein fusion may interfere with the interactions. For instance, we noticed that the colocalization between Atg1 and Atg13 was strong when Atg1 was N-terminally tagged with Pil1-mCherry and used as bait, but was weak when Atg1 was C-terminally tagged with GFP and used as prey (Fig. S1C,D, S2A,B, 4D,E). Considering that in *S. cerevisiae* Atg1 binds Atg13 via its two tandem MIT domains located at the most C-terminus of Atg1 (Fujioka et al., 2014), it is possible that GFP fused at the C-terminus of Atg1 may partially hinder the interaction between Atg1 and Atg13. In our current design of the Pil1 co-tethering assay, bait proteins are always N-terminally tagged and prey proteins can be either N-terminally or C-terminally tagged. Thus, proteins that may be sensitive to tagging at the N-terminus should be C-terminally tagged and used as preys. Third, this assay cannot distinguish whether a protein-protein interaction is direct or bridged by other proteins. Genetically deleting genes encoding proteins that may bridge the interaction, such as other subunits in the same complex, can help address this question. Last, although we have demonstrated that this assay is applicable to transmembrane proteins and nuclear proteins, it remains possible that certain proteins with specific subcellular localizations may be unsuitable for this assay.

In summary, we established a convenient and effective method, the Pil1 co-tethering assay, to allow visual detection of binary, ternary, and quaternary protein-protein interactions in living *S. pombe* cells. For its simplicity and reliability, this method can be used as a routine assay to examine whether two proteins interact, characterize protein-protein interactions in multiprotein complexes, and map interaction regions. It can also be employed to investigate how genetic and environmental changes affect protein-protein interactions. It has the potential to be applied in a large-scale manner if combined with a high-throughput imaging instrument. Even though the Pil1 co-tethering assay is particular suitable for investigating fission yeast proteins in their native cellular context, it can also be used as a heterologous assay system for studying proteins from other organisms.

## Materials and methods

### Fission yeast strains and plasmids

Fission yeast strains used in this study are listed in Table S1, and plasmids used in this study are listed in Table S2. Genetic methods for strain construction and composition of media are as described previously (Forsburg and Rhind, 2006). Deletion strains used in this study were constructed by standard PCR-based gene targeting (Bähler et al., 1998). Plasmids expressing Pil1-mCherry-fused bait proteins or GFP-fused prey proteins under the control of the *41nmt1* (medium-strength *nmt1* promoter) promoter were constructed using modified pDUAL vectors (Matsuyama et al., 2004) (PMID: 24806815). The resulting pDUAL-based plasmids were linearized with NotI digestion and integrated at the *leu1* locus or linearized with MluI digestion and integrated at the *ars1* replication origin region upstream of the *hus5* gene. Plasmids expressing CFP fused Vps34, Atg6, Atg14, or Vps38, and 13Myc fused Vps15 under the control of the *41nmt1* promoter were constructed using the pHIS3H vector (Matsuyama et al., 2008). The resulting pHIS3H-based plasmids were linearized with NotI digestion and integrated at the *his3* locus, except that pHIS3H-P41nmt1-CFP-vps34 was linearized with EcoRV and integrated at the *vps34* locus and pHIS3H-P41nmt1-13Myc-vps15 was linearized with SalI and integrated at the *vps15* locus.

### Fluorescent microscopy

Live-cell imaging was performed using a DeltaVision PersonalDV system (Applied Precision) equipped with an mCherry/YFP/CFP filter set (Chroma 89006 set). Images were acquired with a 100×, 1.4-NA objective using either a Photometrics CoolSNAP HQ2 CCD camera or a Photometrics Evolve 512 EMCCD camera, and analyzed with the SoftWoRx software (GE Healthcare Life Sciences).

### Pil1 co-tethering assay

Proteins analyzed by the Pil1 co-tethering assays were all expressed from plasmids integrated in the genome. Non-integrated episomal plasmids can cause variable expression levels and thus should be avoided. Pil1-mCherry-bait proteins were all expressed under the control of the *41nmt1* promoter. This promoter is strong enough to generate robust fluorescence signal but not too strong to cause abnormal cell morphology and reduced growth rates that can result from strong overexpression of Pil1 (Kabeche et al., 2011). All the prey proteins were expressed from the *41nmt1* promoter except for Atg9-YFP, which was expressed from the *nmt1* promoter. Analyzed strains were cultured to mid-log phase in the EMM medium with appropriate supplements at 30°C. To image the plasma membrane-associated filament-like structures formed by Pil1-mCherry-bait and its interactors, we acquired 5–7 optical Z-sections 0.2 µm apart so that at least in one Z-section the top or bottom plasma membrane was in focus. Then images were processed using the deconvolution algorithm of the SoftWoRx software. The top/bottom Z-section images were shown in most figures. In Fig. 1A and Fig. 3E,G, the mid- plane Z-section images were also shown.

### Computation of the Pearson correlation coefficient (PCC)

The Pearson correlation coefficient (PCC) (Dunn et al., 2011) was used to quantify the degree of colocalization between bait and prey. Imaging data from the corresponding experiments were analyzed using the Coloc 2 plugin of the Fiji distribution of the ImageJ software (http://imagej.net/Coloc_2) (Schindelin et al., 2012). Individual cells in a deconvolved Z-section were outlined and selected as regions of interest (ROIs) using the freehand selection tool. After running the Coloc 2 plugin, Pearson’s R value (no threshold) reported in the ImageJ Log window was recorded for each cell.

### Calculation of buried surface area

The buried surface area in a protein-protein interaction interface was calculated as the sum of the solvent accessible surface areas of the two protein monomers minus the solvent accessible surface area of the complex (Chen et al., 2013). The calculations of solvent accessible surface areas were performed using the website GETAREA (http://curie.utmb.edu/getarea.html) with water represented as a sphere with a radius of 1.4 Å (Fraczkiewicz and Braun, 1998). The individual PDB files submitted to GETAREA were generated by PyMOL based on the solved structure of the budding yeast PtdIns3K complex II (PDB 5DFZ) (Rostislavleva et al., 2015).

### Pho8Δ60 assay in fission yeast

Pho8Δ60 assay was performed as described previously (Yu et al., 2020). Briefly, five OD600 units of cells were harvested and washed with 0.85% NaCl, and then suspended in 200 µl of lysis buffer (20 mM PIPES, pH 6.8, 50 mM KCl, 100 mM KOAc, 10 mM MgSO_4_, 10 µM ZnSO_4_, 0.5% Triton X-100, 2 mM PMSF (freshly added before use)) and incubated at room temperature for 20 min. PMSF was replenished to the final concentration of 4 mM and 0.5- mm-diameter glass beads were added to the samples. Then the cells were disrupted using a FastPrep-24 instrument. After centrifugation, 50 µl of the supernatant was added to 400 µl of reaction buffer (250 mM Tris-HCl, pH 8.5, 10 mM MgSO_4_, 10 µM ZnSO_4_, 0.4% Triton X-100, 5.5 mM 1-naphthyl phosphate disodium salt) to start the reaction. Then samples were incubated at 30°C for 20 min before 500 µl of 1 M glycine-NaOH (pH 11.0) was added to stop the reaction. Fluorescence emission intensity at 472 nm with excitation at 345 nm was measured. Protein concentration was determined by the BCA method.

## Acknowledgments

This work was supported by grants from the Ministry of Science and Technology of the People’s Republic of China and the Beijing municipal government to L.-L.D.

## Abbreviations

Y2H: yeast two-hybrid
FRET: fluorescence resonance energy transfer
BiFC: bimolecular fluorescence complementation
AIM: Atg8-family-interacting motif
NLS: nuclear localization signal

## Supplementary Information

**Figure S1.**
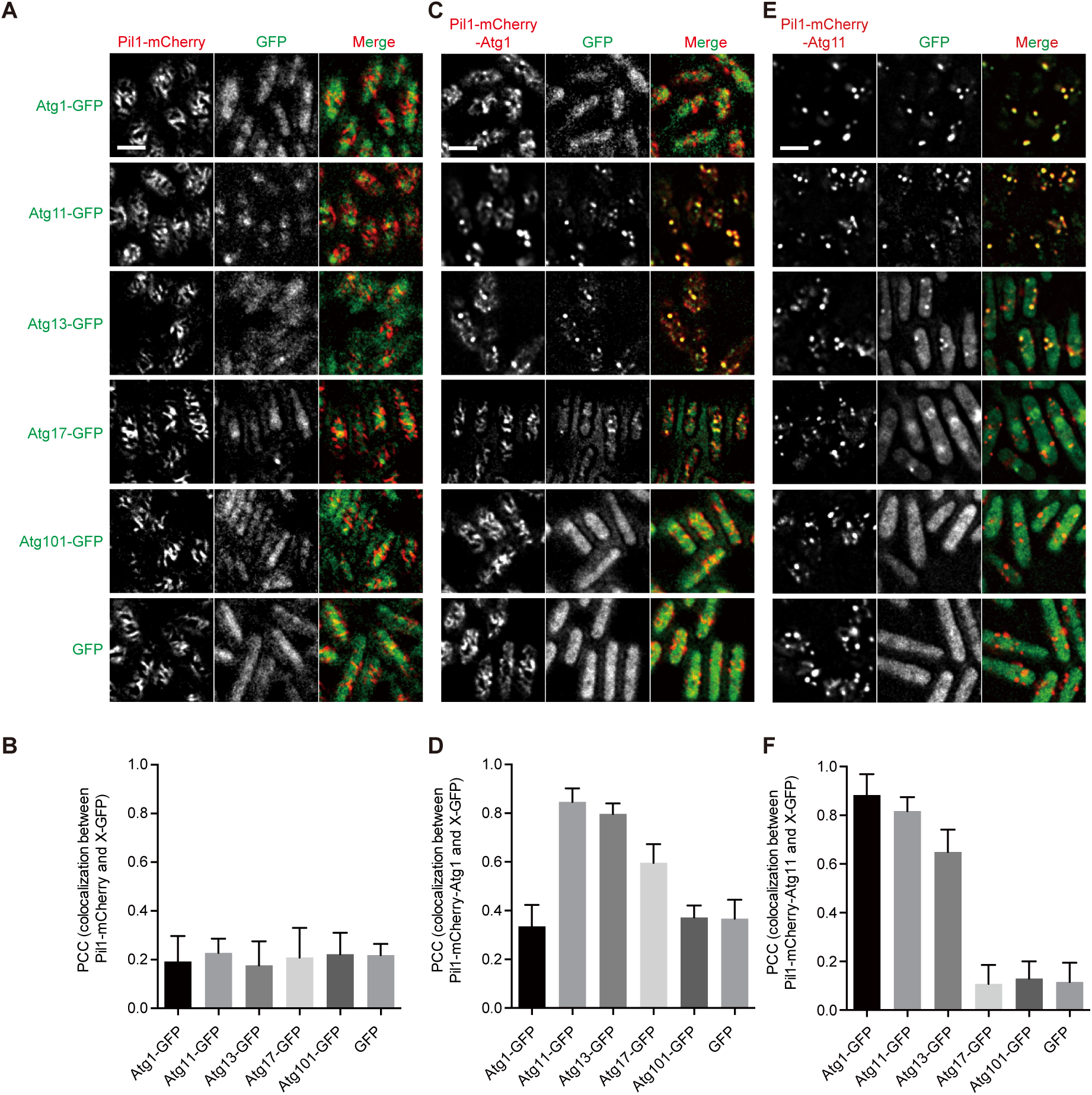
Pil1 co-tethering assay using Atg1 and Atg11 as bait and subunits of the Atg1 complex as prey. (A) Experiments using Pil1-mCherry as a negative control bait. (B) Imaging data from the experiments shown in (A) were analyzed and the PCC values are presented as mean ± s.d. (10 cells). (C) Experiments using the Pil-mCherry-Atg1 bait. (D) Imaging data from the experiments shown in (C) were analyzed and the PCC values are presented as mean ± s.d. (10 cells). (E) Experiments using the Pil-mCherry-Atg11 bait. (F) Imaging data from the experiments shown in (E) were analyzed and the PCC values are presented as mean ± s.d. (10 cells). Scale bars, 5 µm.

**Figure S2.**
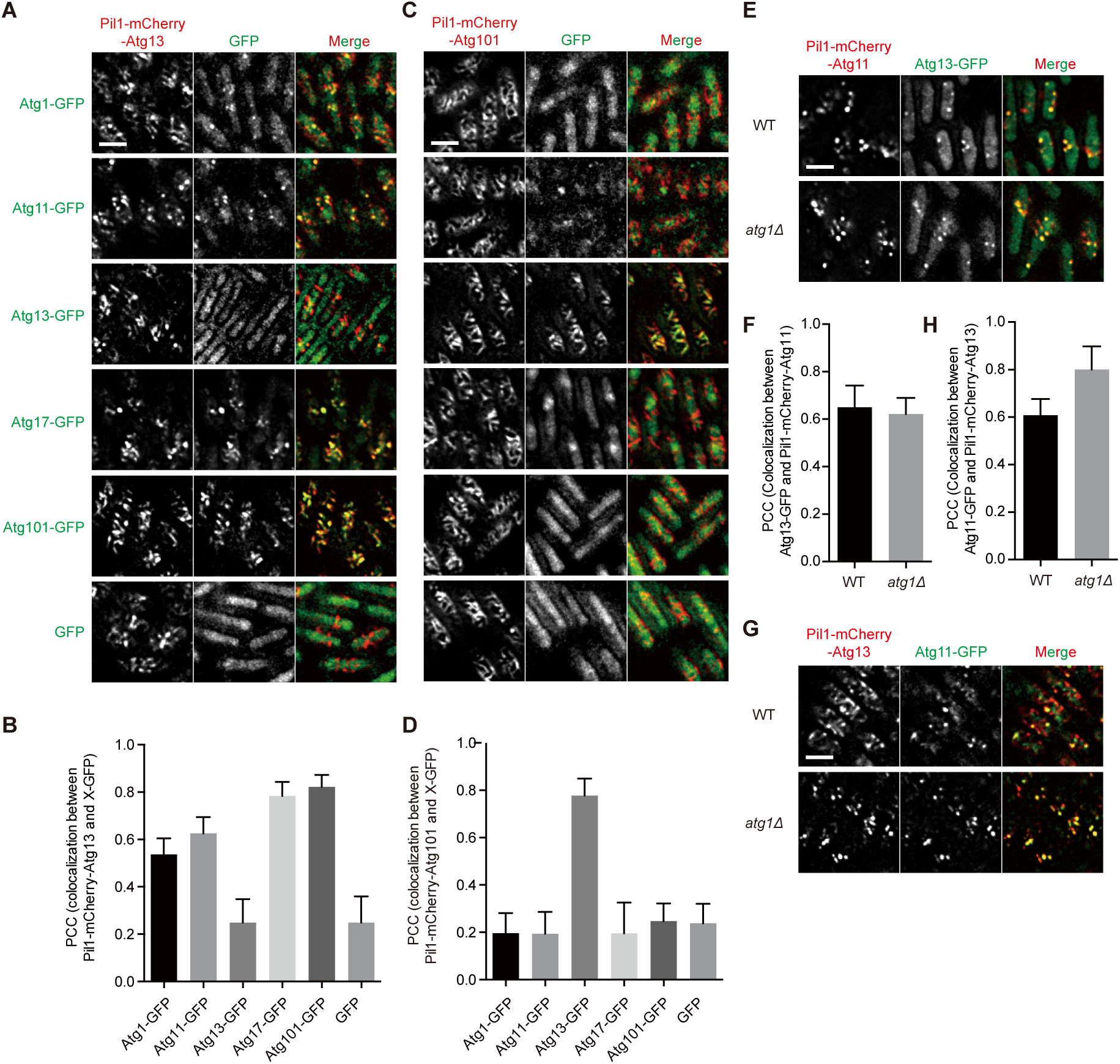
Pil1 co-tethering assay using Atg13 and Atg101 as bait and subunits of the Atg1 complex as prey, and the experiments showing that the interaction between Atg11 and Atg13 is independent of Atg1. (A) Experiments using the Pil-mCherry-Atg13 bait. (B) Imaging data from the experiments shown in (A) were analyzed and the PCC values are presented as mean ± s.d. (10 cells). (C) Experiments using the Pil-mCherry-Atg101 bait. (D) Imaging data from the experiments shown in (C) were analyzed and the PCC values are presented as mean ± s.d. (10 cells). (E) The deletion of *atg1* did not affect the interaction between Atg11 and Atg13 when using Atg11 as bait. (F) Imaging data from the experiments shown in (E) were analyzed and the PCC values are presented as mean ± s.d. (10 cells). (G) The deletion of *atg1* did not affect the interaction between Atg11 and Atg13 when using Atg13 as bait. (H) Imaging data from the experiments shown in (G) were analyzed and the PCC values are presented as mean ± s.d. (10 cells). Scale bars, 5 µm.

**Figure S3.**
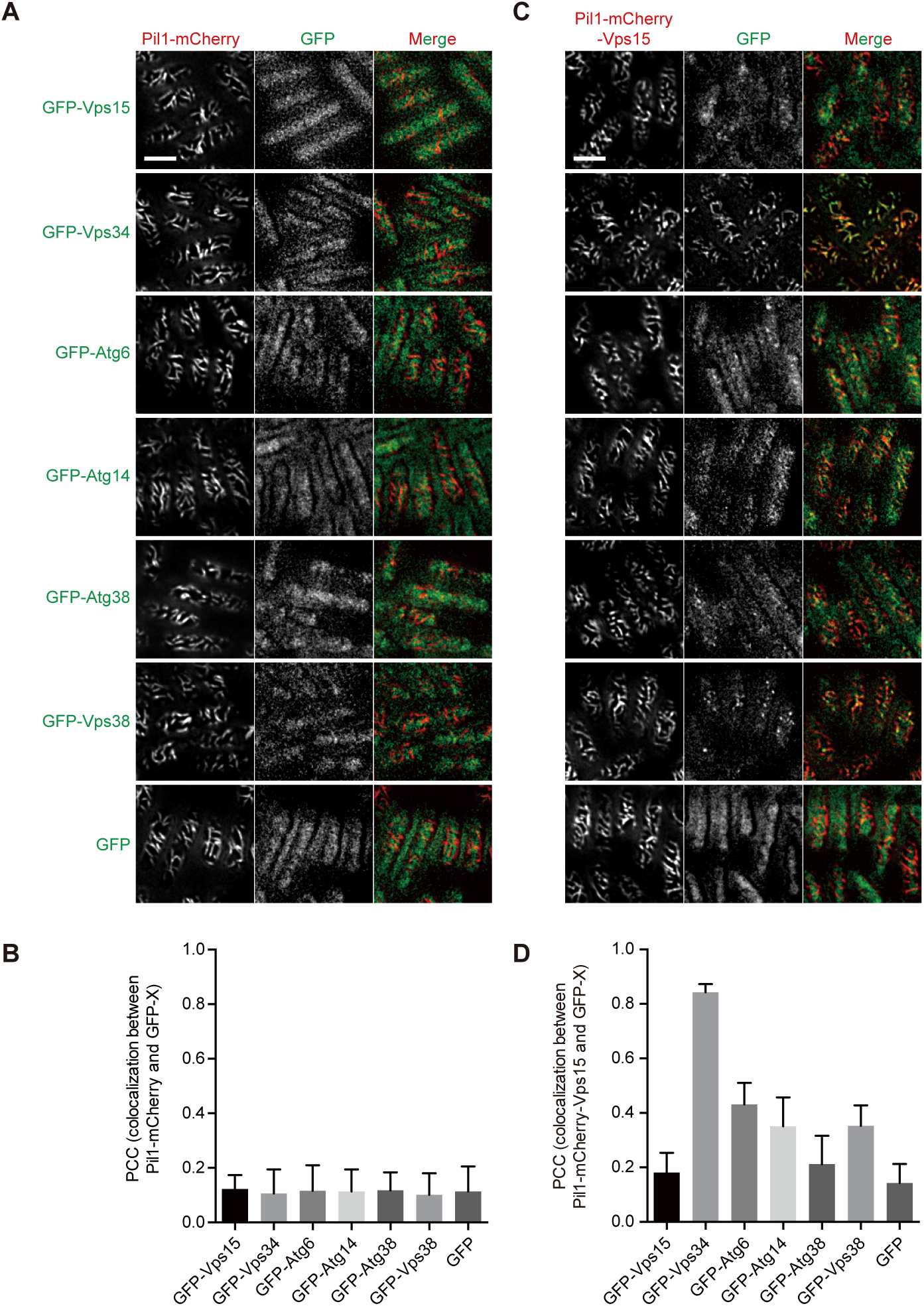
Pil1 co-tethering assay using Vps15 as bait and subunits of PtdIns3K complexes as prey. (A) Experiments using Pil1-mCherry as a negative control bait. (B) Imaging data from the experiments shown in (A) were analyzed and the PCC values are presented as mean ± s.d. (10 cells). (C) Experiments using Pil1-mCherry-Vps15 as bait. (D) Imaging data from the experiments shown in (C) were analyzed and the PCC values are presented as mean ± s.d. (10 cells). Scale bars, 5 µm.

**Figure S4.**
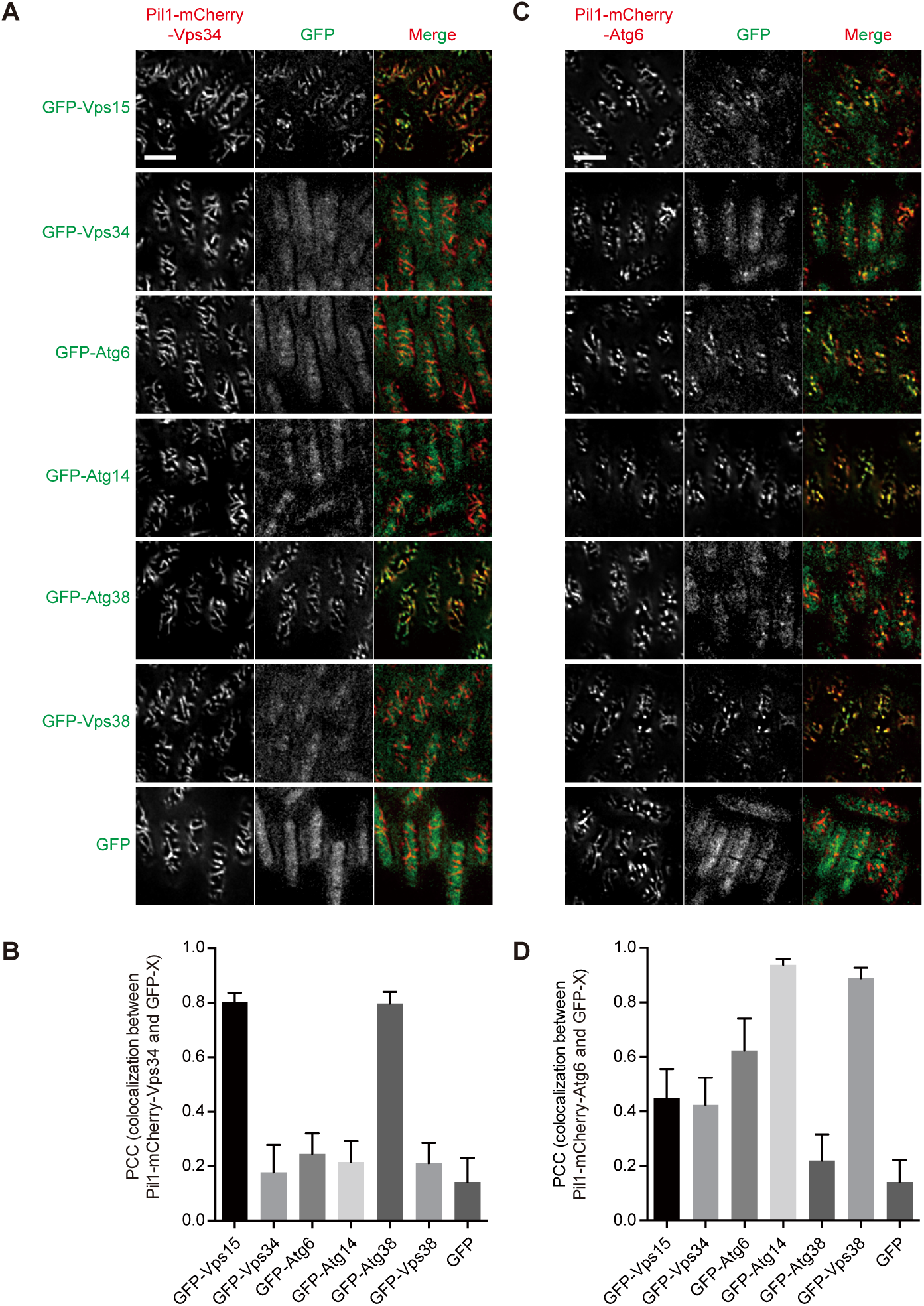
Pil1 co-tethering assay using Vps34 and Atg6 as bait and subunits of PtdIns3K complexes as prey. (A) Experiments using Pil1-mCherry-Vps34 as bait. (B) Imaging data from the experiments shown in (A) were analyzed and the PCC values are presented as mean ± s.d. (10 cells). (C) Experiments using Pil1-mCherry-Atg6 as bait. (D) Imaging data from the experiments shown in (C) were analyzed and the PCC values are presented as mean ± s.d. (10 cells). Scale bars, 5 µm.

**Figure S5.**
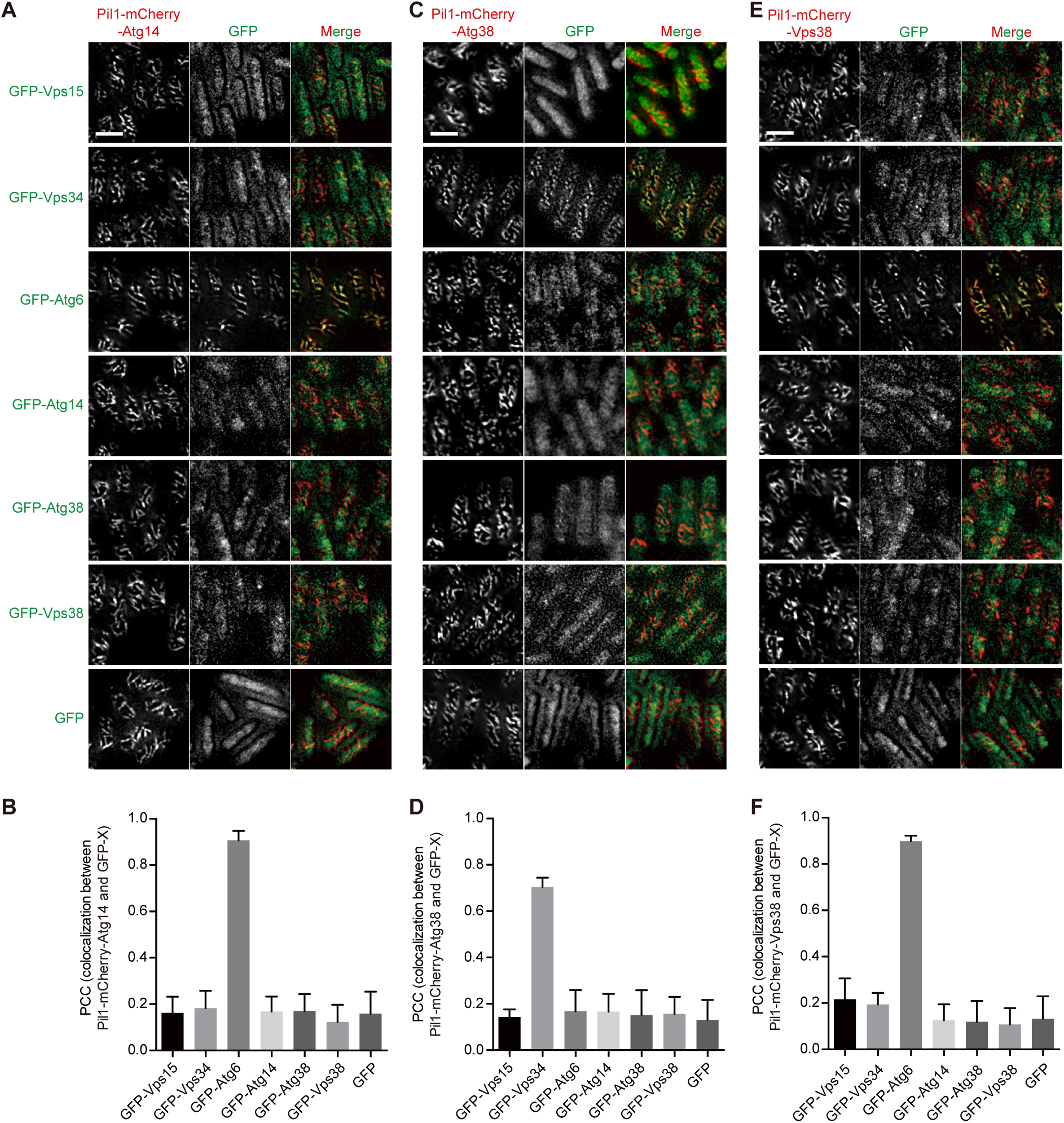
Pil1 co-tethering assay using Atg14, Atg38, and Vps38 as bait and subunits of PtdIns3K complexes as prey. (A) Experiments using Pil1-mCherry-Atg14 as bait. (B) Imaging data from the experiments shown in (A) were analyzed and the PCC values are presented as mean ± s.d. (10 cells). (C) Experiments using Pil1-mCherry-Atg38 as bait. (D) Imaging data from the experiments shown in (C) were analyzed and the PCC values are presented as mean ± s.d. (10 cells). (E) Experiments using Pil1-mCherry-Vps38 as bait. (F) Imaging data from the experiments shown in (E) were analyzed and the PCC values are presented as mean ± s.d. (10 cells). Scale bar, 5 µm.

**Figure S6.**
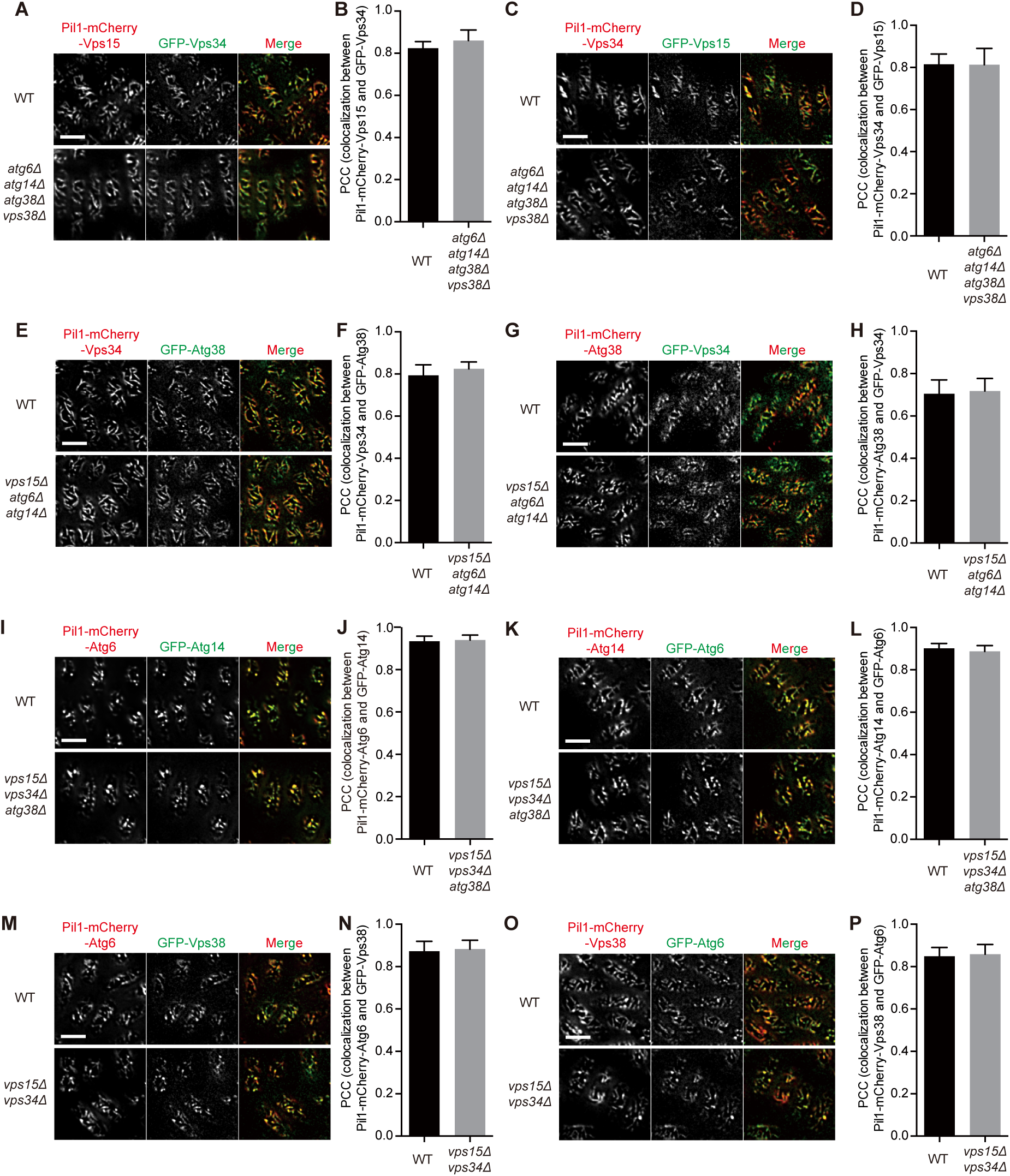
Interactions between Vps15 and Vps34, between Vps34 and Atg38, between Atg6 and Atg14, and between Atg6 and Vps38 are independent of the other subunits of PtdIns3K complexes. (A) Deletion of *atg6*, *atg14*, *atg38*, and *vps38* did not influence the interaction between Vps15 and Vps34 when using Vps15 as bait. (B) Imaging data from the experiments shown in (A) were analyzed and the PCC values are presented as mean ± s.d. (10 cells). (C) Deletion of *atg6*, *atg14*, *atg38*, and *vps38* did not influence the interaction between Vps15 and Vps34 when using Vps34 as bait. (D) Imaging data from the experiments shown in were analyzed and the PCC values are presented as mean ± s.d. (10 cells). (E) Deletion of *vps15*, *atg6*, and *atg14* did not influence the interaction between Vps34 and Atg38 when using Vps34 as bait. (F) Imaging data from the experiments shown in (E) were analyzed and the PCC values are presented as mean ± s.d. (10 cells). (G) Deletion of *vps15*, *atg6*, and *atg14* did not influence the interaction between Vps34 and Atg38 when using Atg38 as bait. (H) Imaging data from the experiments shown in (G) were analyzed and the PCC values are presented as mean ± s.d. (10 cells). (I) Deletion of *vps15*, *vps34*, and *atg38* did not influence the interaction between Atg6 and Atg14 when using Atg6 as bait. (J) Imaging data from the experiments shown in (I) were analyzed and the PCC values are presented as mean ± s.d. (10 cells). (K) Deletion of *vps15*, *vps34*, and *atg38* did not influence the interaction between Atg6 and Atg14 when using Atg14 as bait. (L) Imaging data from the experiments shown in (K) were analyzed and the PCC values are presented as mean ± s.d. (10 cells). (M) Deletion of *vps15* and *vps34* did not influence the interaction between Atg6 and Vps38 when using Atg6 as bait. (N) Imaging data from the experiments shown in (M) were analyzed and the PCC values are presented as mean ± s.d. (10 cells). (O) Deletion of *vps15* and *vps34* did not influence the interaction between Atg6 and Vps38 when using Vps38 as bait. (P) Imaging data from the experiments shown in (O) were analyzed and the PCC values are presented as mean ± s.d. (10 cells). Scale bars, 5 µm.

**Figure S7.**
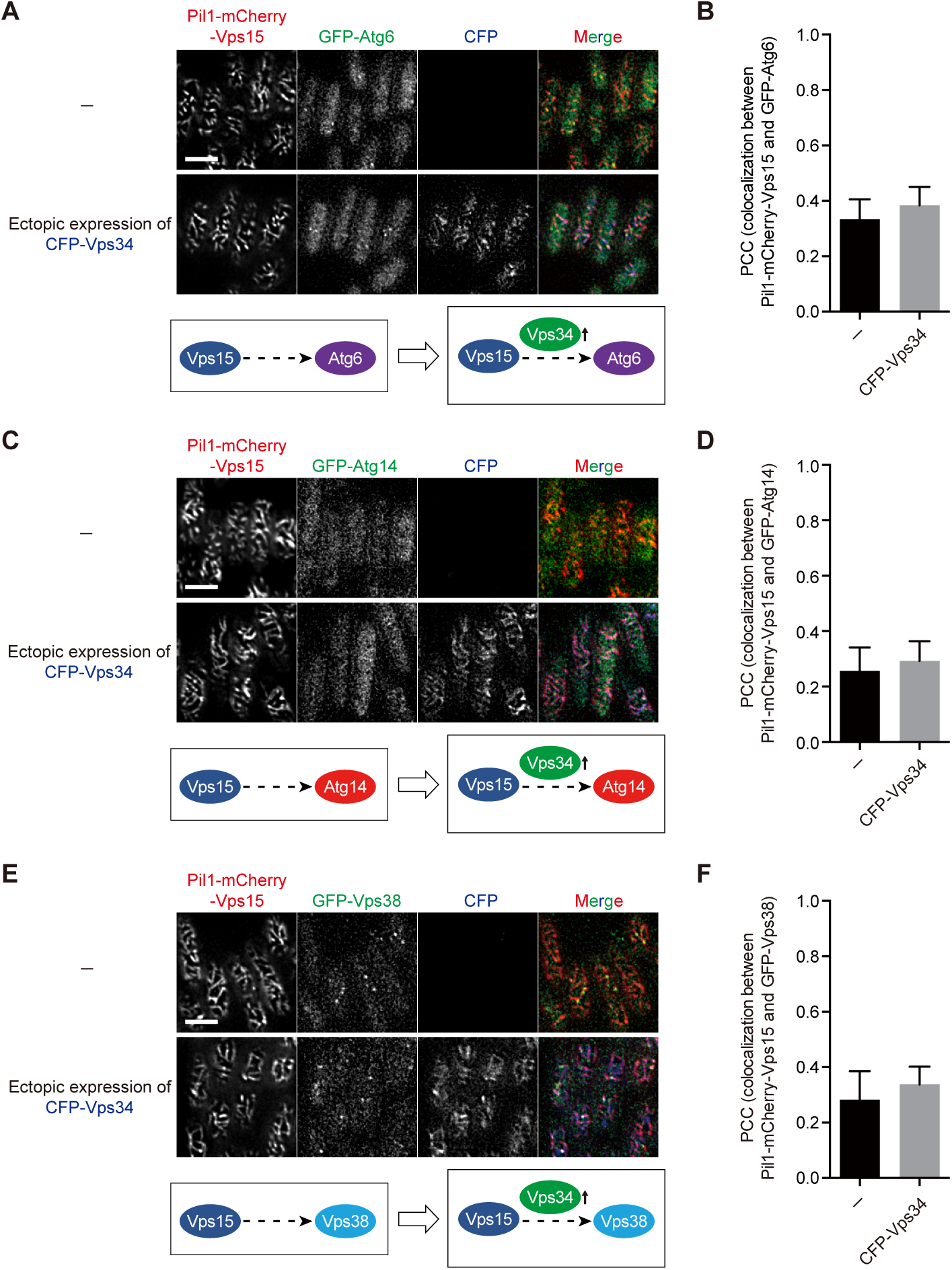
Atg6, Atg14, and Vps38 individually does not interact with the Vps15-Vps34 subcomplex. (A) Ectopic expression of Vps34 did not lead to the colocalization of Vps15 and Atg6. (B) Imaging data from the experiments shown in (A) were analyzed and the PCC values are presented as mean ± s.d. (10 cells). (C) Ectopic expression of Vps34 did not lead to the colocalization of Vps15 and Atg14. (D) Imaging data from the experiments shown in (C) were analyzed and the PCC values are presented as mean ± s.d. (10 cells). (E) Ectopic expression of Vps34 did not lead to the colocalization of Vps15 and Vps38. (F) Imaging data from the experiments shown in (E) were analyzed and the PCC values are presented as mean ± s.d. (10 cells). Scale bars, 5 µm.

**Figure S8.**
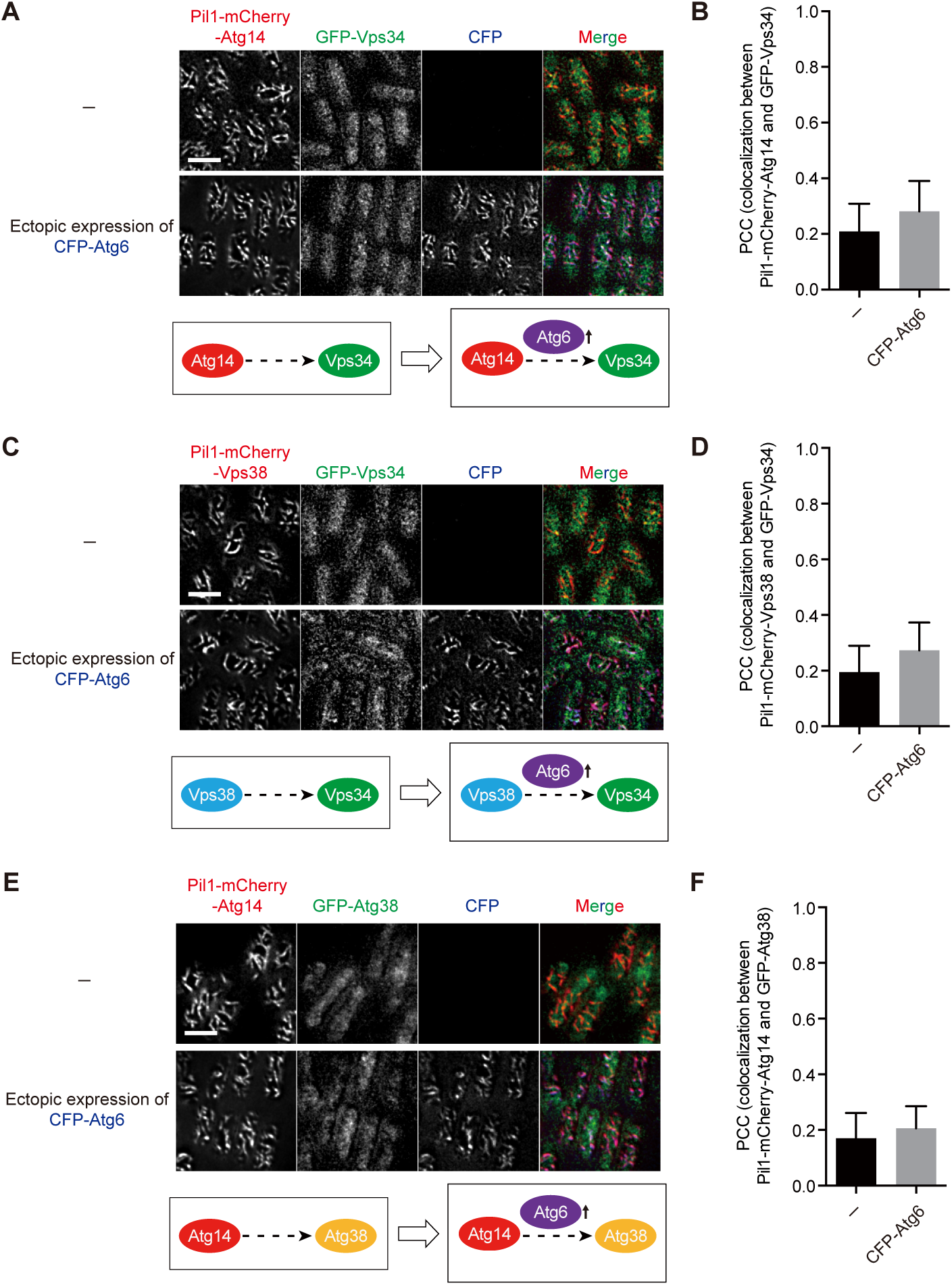
The Atg14-Atg6 subcomplex and the Vps38-Atg6 subcomplex do not interact with Vps34, and the Atg14-Atg6 subcomplex does not interact with Atg38. (A) Ectopic expression of Atg6 did not lead to the colocalization of Atg14 and Vps34. (B) Imaging data from the experiments shown in (A) were analyzed and the PCC values are presented as mean ± s.d. (10 cells). (C) Ectopic expression of Atg6 did not lead to the colocalization of Vps38 and Vps34. (D) Imaging data from the experiments shown in (C) were analyzed and the PCC values are presented as mean ± s.d. (10 cells). (E) Ectopic expression of Atg6 did not lead to the colocalization of Atg14 and Atg38. (F) Imaging data from the experiments shown in (E) were analyzed and the PCC values are presented as mean ± s.d. (10 cells). Scale bars, 5 µm.

**Table S1.**
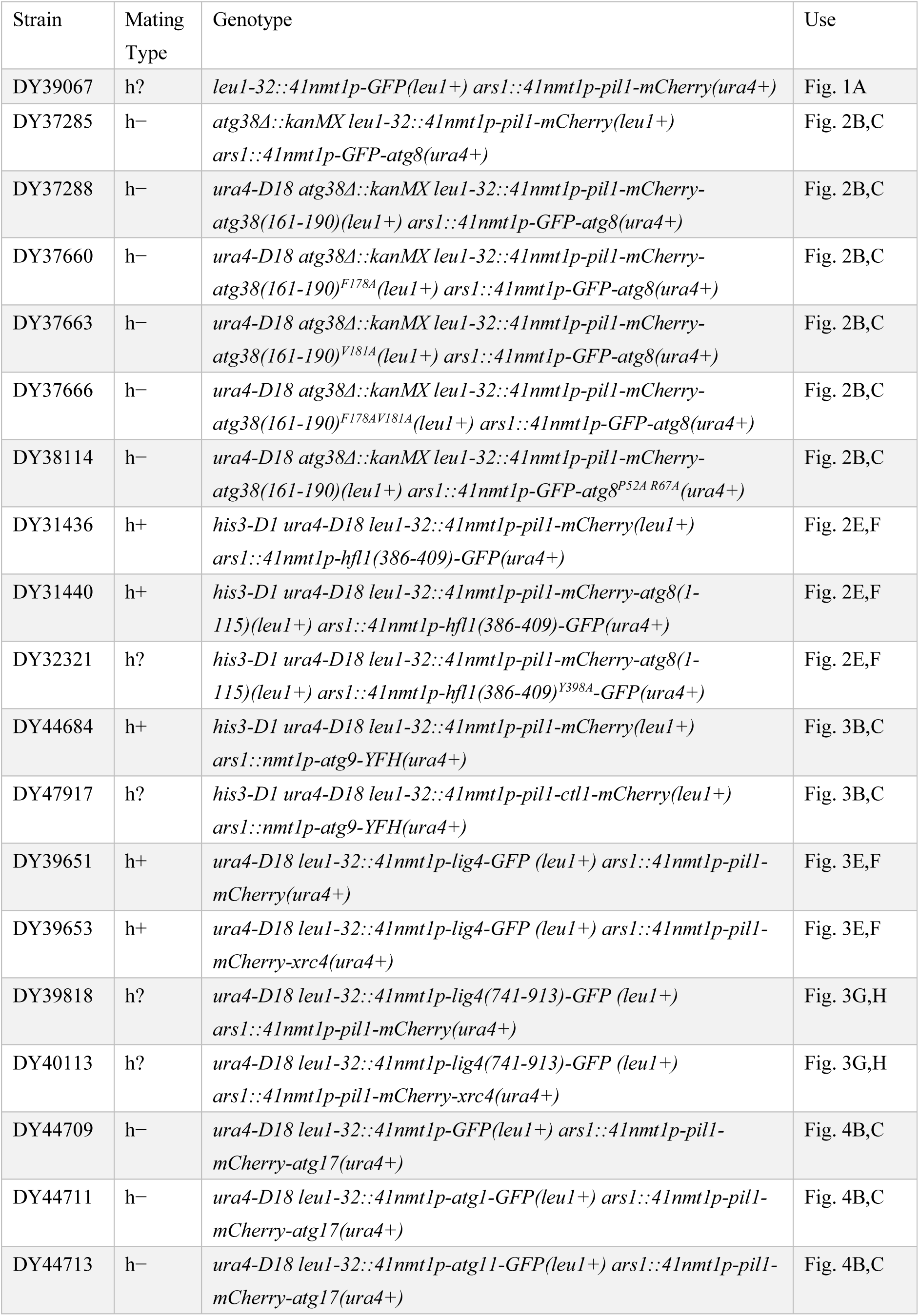
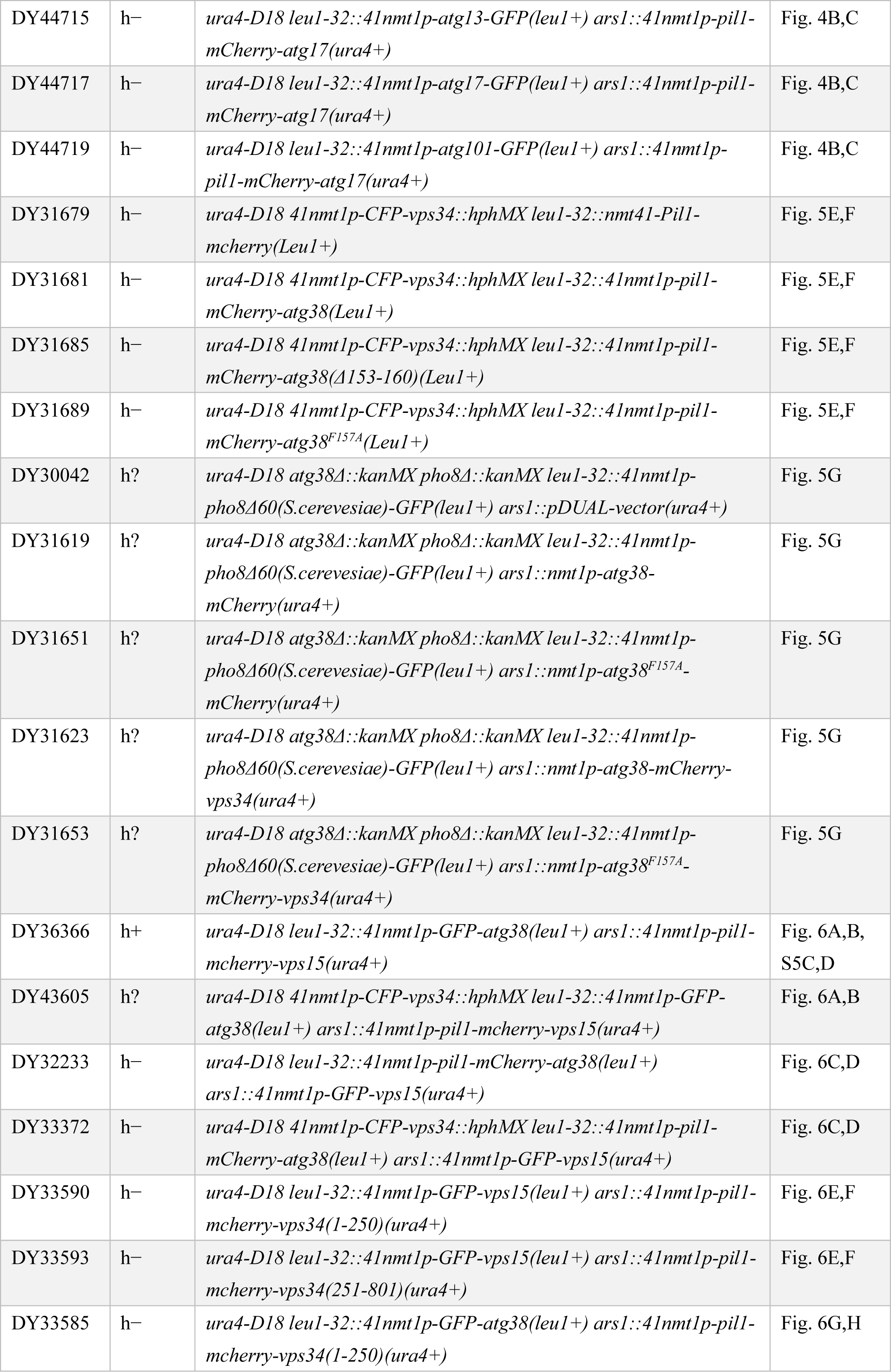
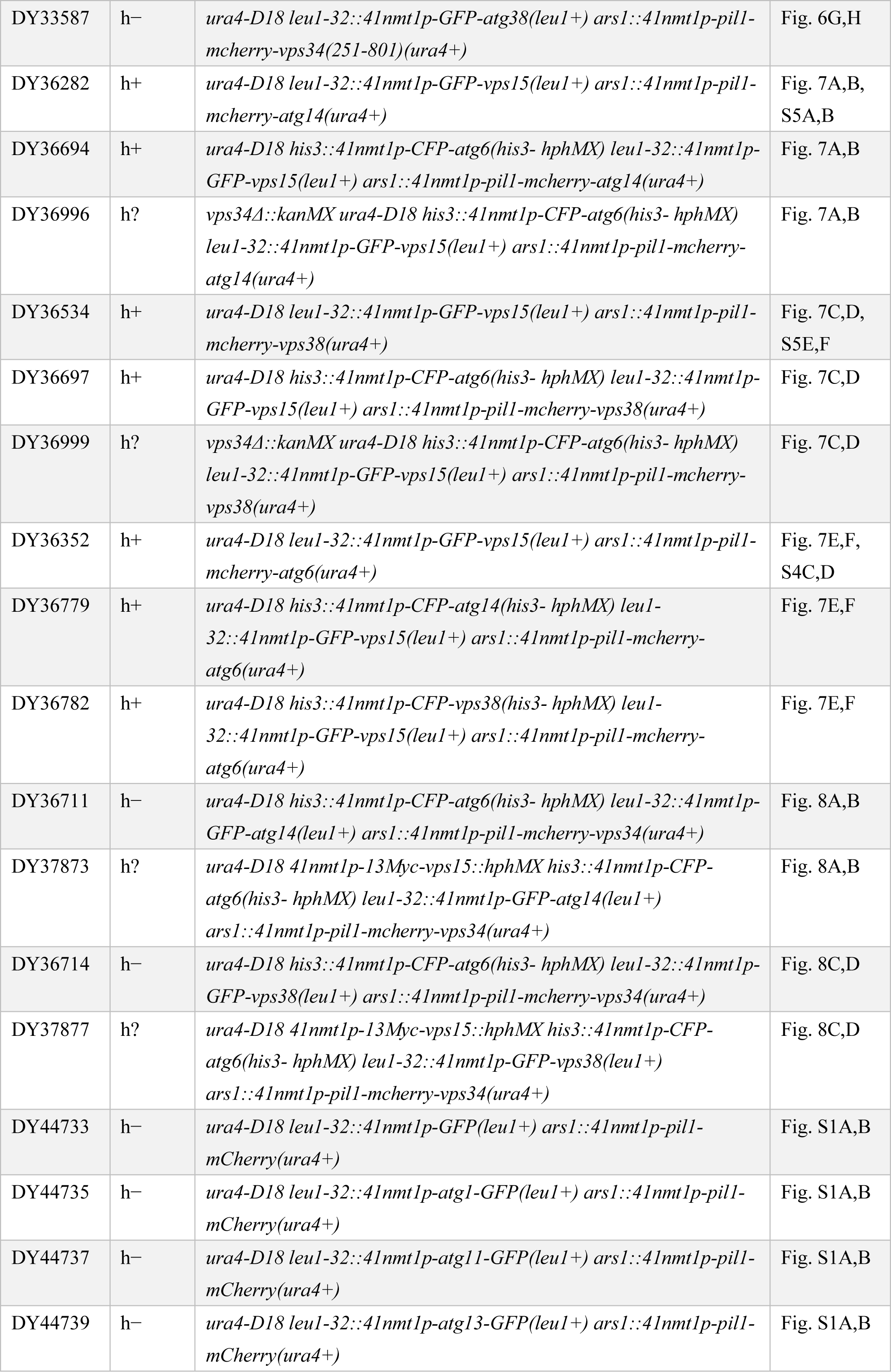
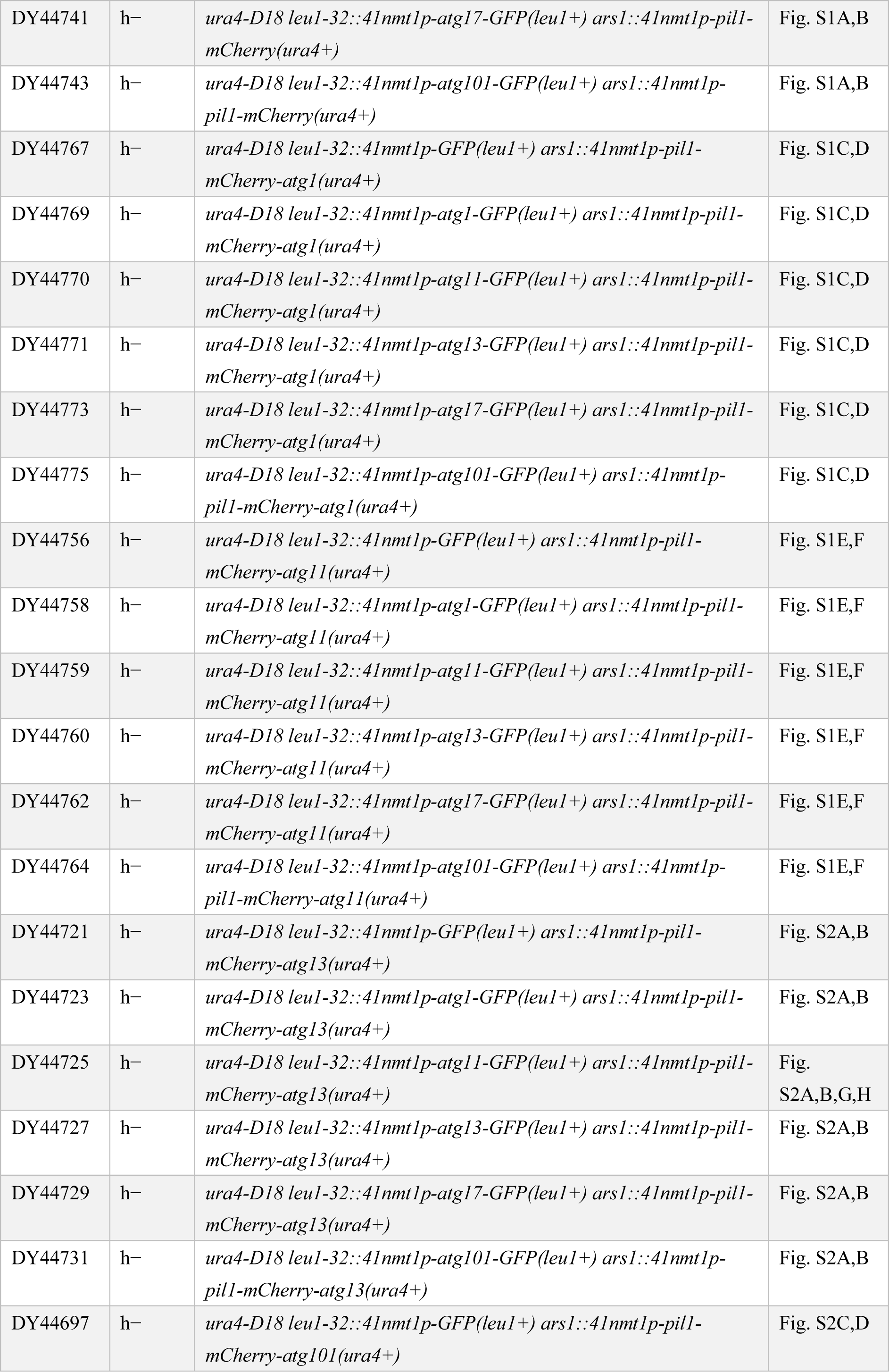
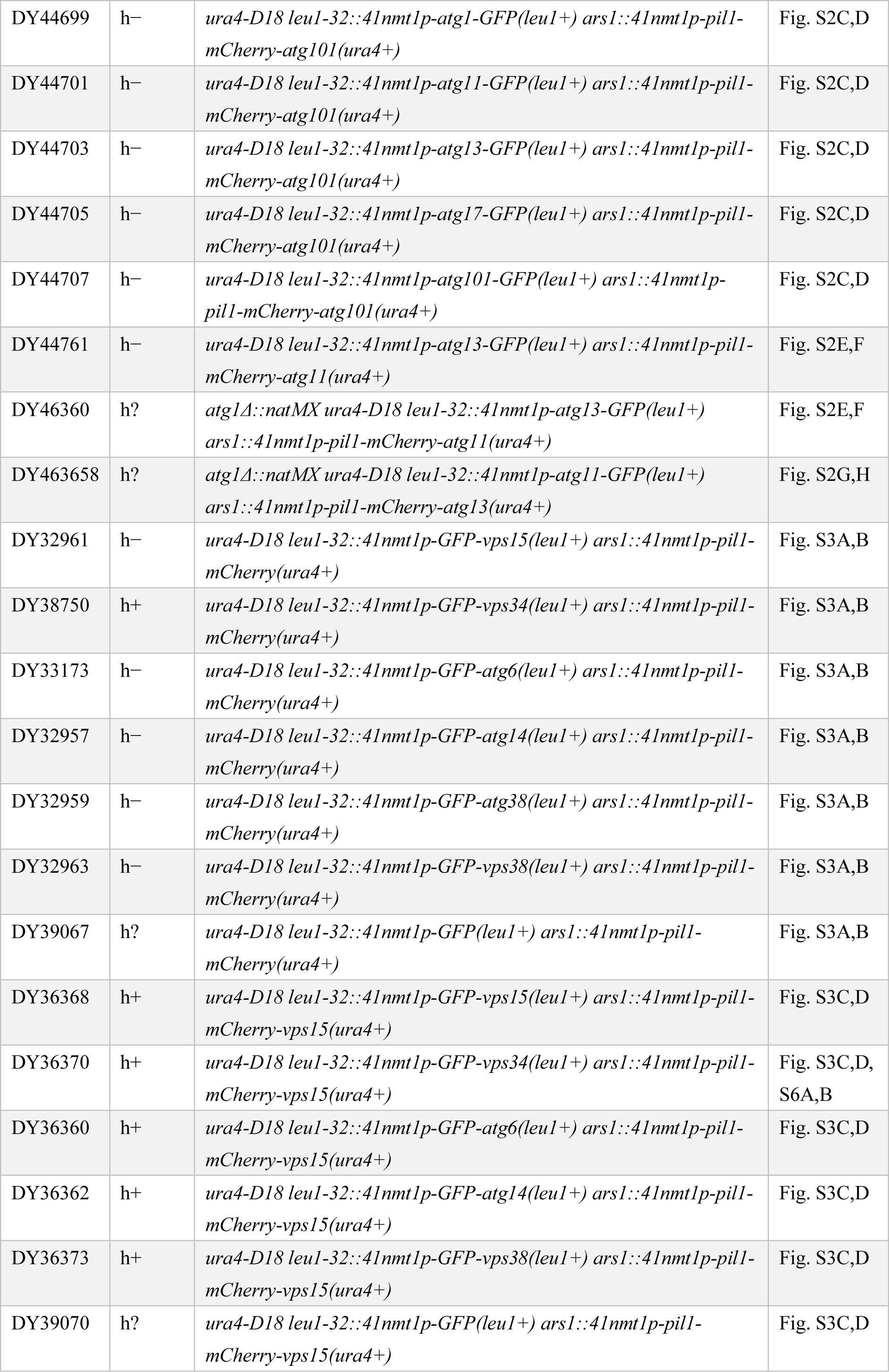
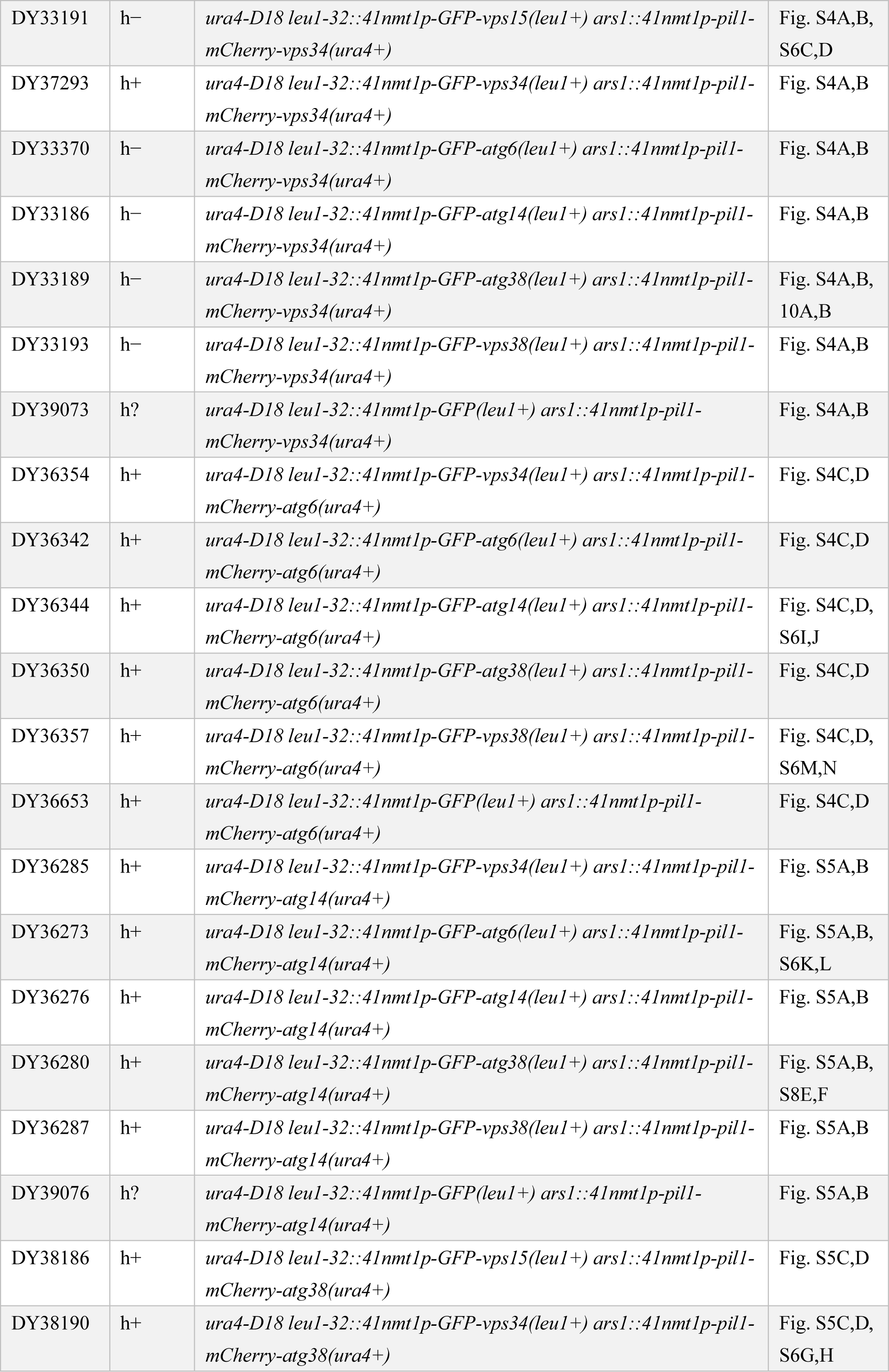
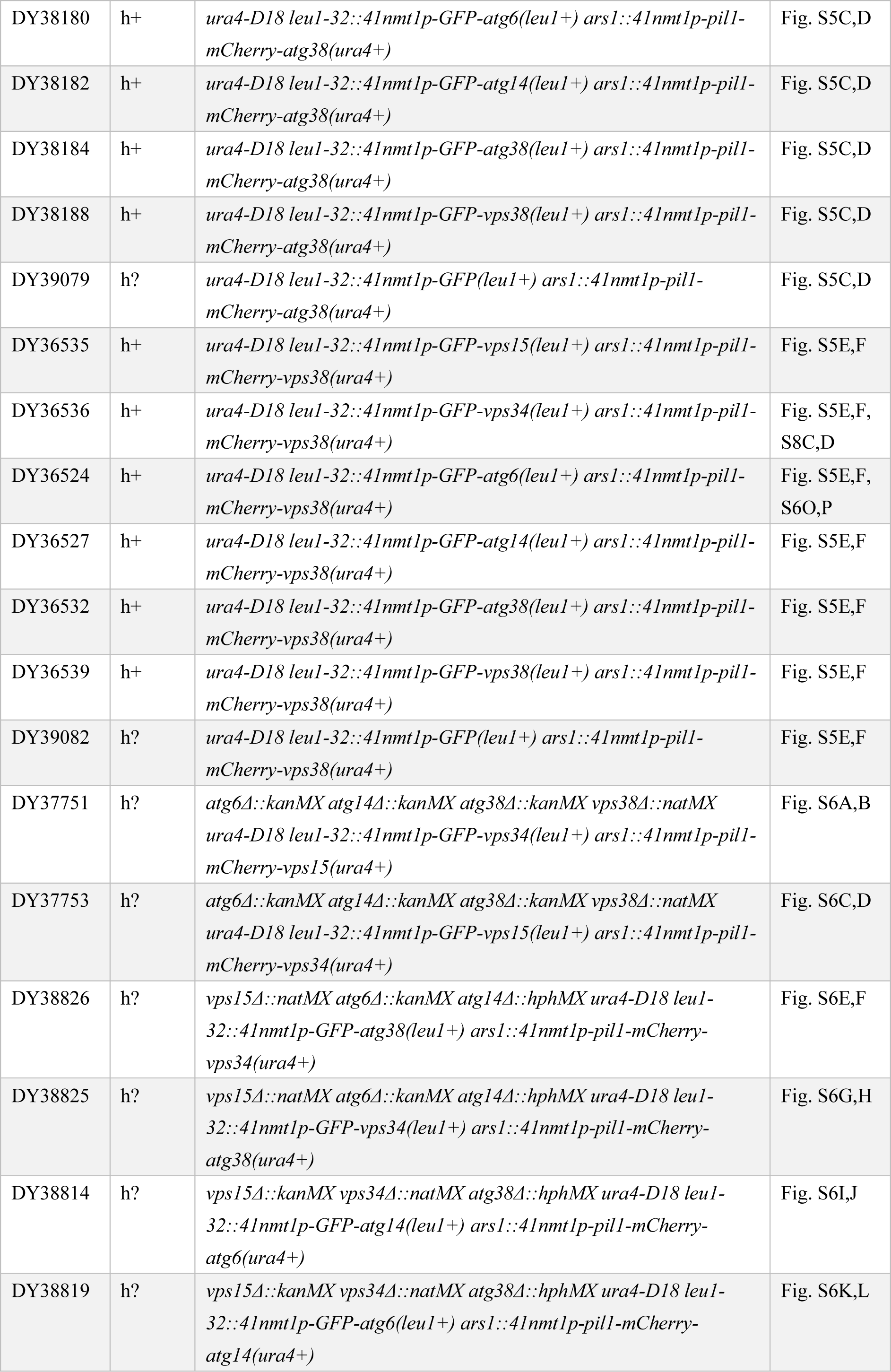
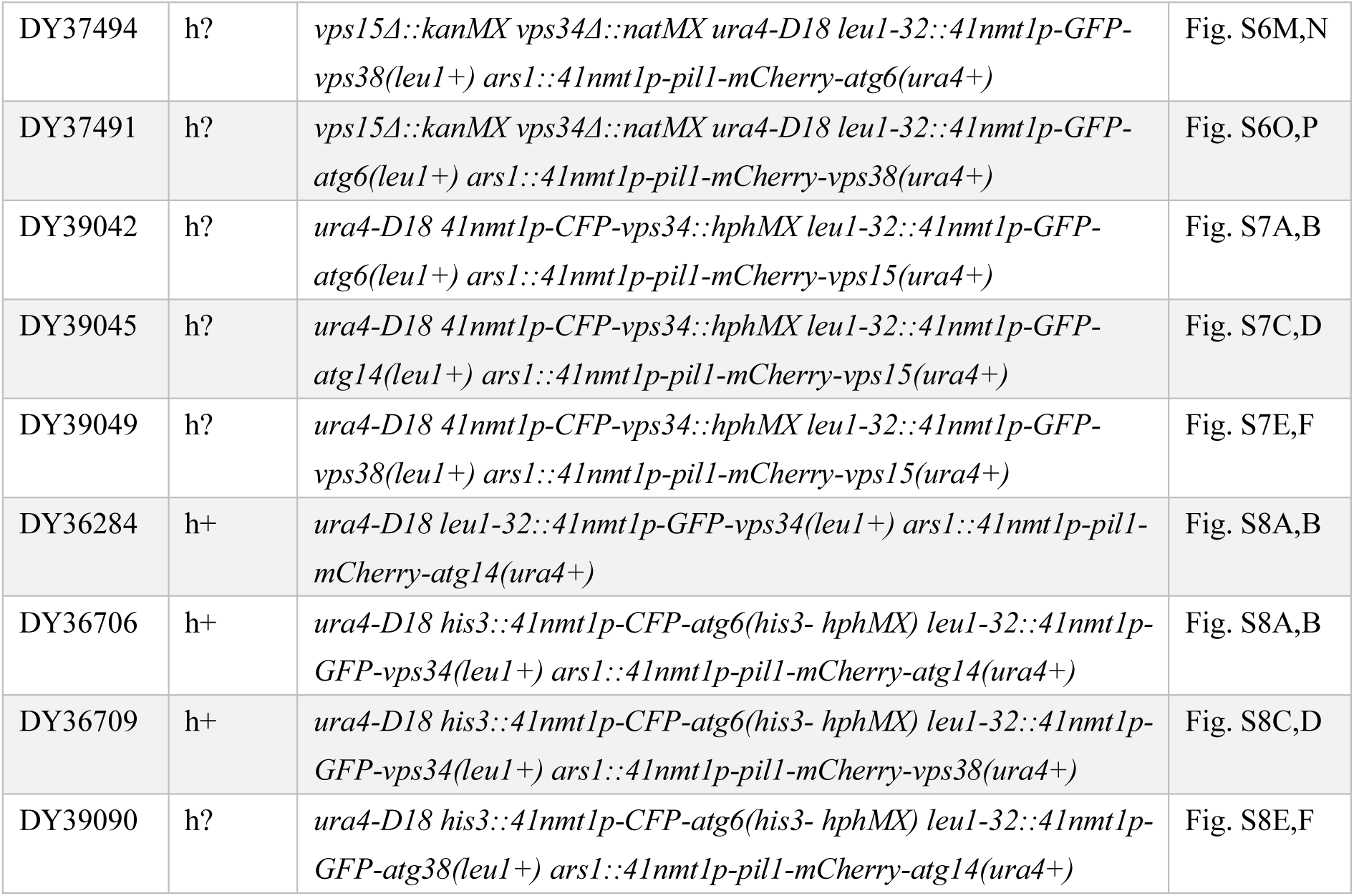
Fission yeast strains used in this study

**Table S2.** Plasmids used in this study

| Name | Descriptive name | Description |
| --- | --- | --- |
| pDB4924 | pDUAL-*4lnmt1p*-Pil1-mCherry | pDUAL plasmid expressing Pil1-mCherry from *4lnmt1* promoter |
| pDB4925 | pDUAL-*4lnmt1p*-Pil1-mCherry-Atg38(161-190) | pDUAL plasmid expressing Pil1-mCherry-Atg38(161-190) from *4lnmt1* promoter |
| pDB4926 | pDUAL-*4lnmt1p*-Pil1-mCherry-Atg38(161-190)<sup>F178A</sup> | pDUAL plasmid expressing Pil1-mCherry-Atg38(161-190)<sup>F178A</sup> from *4lnmt1* promoter |
| pDB4927 | pDUAL-*4lnmt1p*-Pil1-mCherry-Atg38(161-190)<sup>V181A</sup> | pDUAL plasmid expressing Pil1-mCherry-Atg38(161-190)<sup>V181A</sup> from *4lnmt1* promoter |
| pDB4928 | pDUAL-*4lnmt1p*-Pil1-mCherry-Atg38(161-190)<sup>F178A V181A</sup> | pDUAL plasmid expressing Pil1-mCherry-Atg38(161-190)<sup>F178A V181A</sup> from *4lnmt1* promoter |
| pDB4658 | pDUAL-*4lnmt1p*-GFP-Atg8 | pDUAL plasmid expressing GFP-Atg8 from *4lnmt1* promoter |
| pDB4659 | pDUAL-*4lnmt1p*-GFP-Atg8<sup>P52A R67A</sup> | pDUAL plasmid expressing GFP-Atg8<sup>P52A R67A</sup> from *4lnmt1* promoter |
| pDB4929 | pDUAL-*4lnmt1p*-Pil1-mCherry-Atg8(1-115) | pDUAL plasmid expressing Pil1-mCherry-Atg8(1-115) from *4lnmt1* promoter |
| pDB4923 | pDUAL-*4lnmt1p*-Pil1-mCherry-Ctl1 | pDUAL plasmid expressing Pil1-mCherry-Ctl1 from *4lnmt1* promoter |
| pDB4922 | pDUAL-*nmt1p*-Atg9-YFH | pDUAL plasmid expressing Atg9-YFH from *nmt1* promoter |
| pDB4930 | pDUAL-*4lnmt1p*-Pil1-mCherry-Xrc4 | pDUAL plasmid expressing Pil1-mCherry-Xrc4 from *4lnmt1* promoter |
| pDB4931 | pDUAL-*4lnmt1p*-Lig4-GFP | pDUAL plasmid expressing Lig4-GFP from *4lnmt1* promoter |
| pDB4932 | pDUAL-*4lnmt1p*-Lig4(741-913)-GFP | pDUAL plasmid expressing Lig4(741-913)-GFP from *4lnmt1* promoter |
| pDB4933 | pDUAL-*4lnmt1p*-Pil1-mCherry-Atg1 | pDUAL plasmid expressing Pil1-mCherry-Atg1 from *4lnmt1* promoter |
| pDB4934 | pDUAL-*4lnmt1p*-Pil1-mCherry-Atg11 | pDUAL plasmid expressing Pil1-mCherry-Atg11 from *4lnmt1* promoter |
| pDB4935 | pDUAL-*4lnmt1p*-Pil1-mCherry-Atg13 | pDUAL plasmid expressing Pil1-mCherry-Atg13 from *4lnmt1* promoter |
| pDB4936 | pDUAL-*4lnmt1p*-Pil1-mCherry-Atg17 | pDUAL plasmid expressing Pil1-mCherry-Atg17 from *4lnmt1* promoter |
| pDB4937 | pDUAL-*4lnmt1p*-Pil1-mCherry-Atg101 | pDUAL plasmid expressing Pil1-mCherry-Atg101 from *4lnmt1* promoter |
| pDB4938 | pDUAL-*4lnmt1p*-Atg1-GFP | pDUAL plasmid expressing Atg1-GFP from *4lnmt1* promoter |
| pDB4939 | pDUAL-*4lnmt1p*-Atg11-GFP | pDUAL plasmid expressing Atg11-GFP from *4lnmt1* promoter |
| pDB4940 | pDUAL-*4lnmt1p*-Atg13-GFP | pDUAL plasmid expressing Atg13-GFP from *4lnmt1* promoter |
| pDB4941 | pDUAL-*4lnmt1p*-Atg17-GFP | pDUAL plasmid expressing Atg17-GFP from *4lnmt1* promoter |
| pDB4942 | pDUAL-*4lnmt1p*-Atg101-GFP | pDUAL plasmid expressing Atg101-GFP from *4lnmt1* promoter |
| pDB4943 | pDUAL-*4lnmt1p*-Pil1-mCherry-Vps15 | pDUAL plasmid expressing Pil1-mCherry-Vps15 from *4lnmt1* promoter |
| pDB4944 | pDUAL-*4lnmt1p*-Pil1-mCherry-Vps34 | pDUAL plasmid expressing Pil1-mCherry-Vps34 from *4lnmt1* promoter |
| pDB4945 | pDUAL-*4lnmt1p*-Pil1-mCherry-Atg6 | pDUAL plasmid expressing Pil1-mCherry-Atg6 from *4lnmt1* promoter |
| pDB4946 | pDUAL-*4lnmt1p*-Pil1-mCherry-Atg14 | pDUAL plasmid expressing Pil1-mCherry-Atg14 from *4lnmt1* promoter |
| pDB4947 | pDUAL-*4lnmt1p*-Pil1-mCherry-Atg38 | pDUAL plasmid expressing Pil1-mCherry-Atg38 from *4lnmt1* promoter |
| pDB4948 | pDUAL-*4lnmt1p*-Pil1-mCherry-Vps38 | pDUAL plasmid expressing Pil1-mCherry-Vps38 from *4lnmt1* promoter |
| pDB4949 | pDUAL-*4lnmt1p*-Pil1-mCherry-Vps34(1-250) | pDUAL plasmid expressing Pil1-mCherry-Vps34(1-250) from *4lnmt1* promoter |
| pDB4950 | pDUAL-*4lnmt1p*-Pil1-mCherry-Vps34(251-801) | pDUAL plasmid expressing Pil1-mCherry-Vps34(251-801) from *4lnmt1* promoter |
| pDB4951 | pDUAL-*4lnmt1p*-GFP-Vps15 | pDUAL plasmid expressing GFP-Vps15 from *4lnmt1* promoter |
| pDB4952 | pDUAL-*4lnmt1p*-GFP-Vps34 | pDUAL plasmid expressing GFP-Vps34 from *4lnmt1* promoter |
| pDB4953 | pDUAL-*4lnmt1p*-GFP-Atg6 | pDUAL plasmid expressing GFP-Atg6 from *4lnmt1* promoter |
| pDB4954 | pDUAL-*4lnmt1p*-GFP-Atg14 | pDUAL plasmid expressing GFP-Atg14 from *4lnmt1* promoter |
| pDB4955 | pDUAL-*4lnmt1p*-GFP-Atg38 | pDUAL plasmid expressing GFP-Atg38 from *4lnmt1* promoter |
| pDB4956 | pDUAL-*4lnmt1p*-GFP-Vps38 | pDUAL plasmid expressing GFP-Vps38 from *4lnmt1* promoter |
| pDB4957 | pHIS3H-*4lnmt1p*-CFP-Vps34 | pHIS3H plasmid expressing CFP-Vps34 from *4lnmt1* promoter |
| pDB4958 | pHIS3H-*4lnmt1p*-CFP-Atg6 | pHIS3H plasmid expressing CFP-Atg6 from *4lnmt1* promoter |
| pDB4959 | pHIS3H-*4lnmt1p*-CFP-Atg14 | pHIS3H plasmid expressing CFP-Atg14 from *4lnmt1* promoter |
| pDB4960 | pHIS3H-*4lnmt1p*-CFP-Vps38 | pHIS3H plasmid expressing CFP-Vps38 from *4lnmt1* promoter |
| pDB4961 | pHIS3H-*4lnmt1p*-13Myc-Vps15 | pHIS3H plasmid expressing 13Myc-Vps15 from *4lnmt1* promoter |
| pDB4972 | pDUAL-*nmt1p*-Atg38-mCherry | pDUAL plasmid expressing Atg38-mCherry from *nmt1* promoter |
| pDB4973 | pDUAL-*nmt1p*-Atg38<sup>F157A</sup>-mCherry | pDUAL plasmid expressing Atg38<sup>F157A</sup>-mCherry from *nmt1* promoter |
| pDB4974 | pDUAL-*nmt1p*-Atg38-mCherry-Vps34 | pDUAL plasmid expressing Atg38-mCherry-Vps34 from *nmt1* promoter |
| pDB4975 | pDUAL-*nmt1p*-Atg38<sup>F157A</sup>-mcherry-vps34 | pDUAL plasmid expressing Atg38<sup>F157A</sup>-mCherry-Vps34 from *nmt1* promoter |

